# Theta cycle dynamics of spatial representations in the lateral septum

**DOI:** 10.1101/2023.07.17.549368

**Authors:** Katarzyna Bzymek, Fabian Kloosterman

## Abstract

An internal representation of the environment – or map – allows animals to evaluate multiple routes and adapt their navigation strategy to current needs and future goals. The hippocampal formation plays a crucial role in learning a spatial map and using the map for goal-directed navigation. The lateral septum forms a major node for connections between the hippocampus and subcortical brain regions that could link the spatial map to motivation and reward processing centers such as the ventral tegmental area and hypothalamus. It is not known, however, how the lateral septum contributes to the processing of spatial information and route planning.

In this study, we investigated the temporal dynamics of spatial representations in the lateral septum. Neuropixels probes were used to record cellular activity along the dorsal-ventral extent of the lateral septum while rats performed one of two spatial navigation tasks in a Y-maze. The activity of a large fraction of cells was theta rhythmic and a subset of cells showed evidence of being active on alternate theta cycles (theta cycle skipping). Both theta rhythmicity and cycle skipping were strongest in the dorsal lateral septum. Similarly, spatially selective firing was most prominent in the dorsal lateral septum. Using neural decoding, we show that the lateral septum cell population encodes both the current location and alternatingly the possible future paths within single theta cycles when rats approach the choice point in the maze.

Our data further shows that the alternating expression of spatial representations in the lateral septum is task-dependent, such that it is strongest when the task also requires the animals to alternate between rewarded goal arms. These data suggest that task demands and experience shape which representations are activated near a choice point. The lateral septum receives strong input from hippocampal place cells, and while there may be integration and transformation of incoming spatial signals, our findings support the idea that hippocampal spatial representations and their temporal dynamics are conveyed to subcortical projection areas through the lateral septum.

## Introduction

The ability to navigate the environment in search for food is crucial for the survival of mammals. An internal representation of the environment – or map – allows animals to evaluate multiple routes and adapt their navigation strategy to current needs and future goals. The hippocampal formation plays a crucial role in learning a spatial map and using the map for goal-directed navigation (Moser et al., 2017). Through its major projections to the lateral septum (Risold and Swanson, 1997), the hippocampus modulates subcortical networks that control motivational, reward, and social aspects of spatial behavior (Besnard and Leroy, 2022). However, how spatial signals from the hippocampus are translated into lateral septum output is not well understood.

Hippocampal cell populations represent current locations and upcoming trajectories through spatially selective firing and place cell sequences (O’Keefe, 1976; Wilson and McNaughton, 1993). These hippocampal spatial representations are expressed cyclically during ∼8 Hz theta oscillations (Colgin, 2013; Dragoi and Buzsáki, 2006; Foster and Wilson, 2007). In each theta cycle, sensory input and the hippocampal attractor network interact to generate short “sweeps” of place cell activity along the path ahead or behind the current location in the environment (Johnson and Redish, 2007; Wang et al., 2020). When two paths intersect, the place cell representations of the two possible future paths are expressed in alternating fashion (Kay et al., 2020), possibly reflecting a competition between two attractor states. At the single-cell level, this phenomenon is observed as spiking activity that “skips” theta cycles (i.e., enhanced rhythmicity at half theta frequency). Theta cycle skipping and the associated theta sweeps have been linked to the deliberation about the upcoming choice (Johnson and Redish, 2007; Robinson and Brandon, 2021).

There is evidence that theta cycle skipping is present in cortical and thalamic areas with connections to the hippocampus (Brandon et al., 2013; Jankowski et al., 2014; Tang et al., 2021). The question addressed here is if and how theta cycle skipping dynamics of hippocampal spatial representations are relayed to and transformed in the lateral septum. Available evidence indicate that the activity of lateral septum neurons is modulated by the hippocampal theta rhythm (Bender et al., 2015; Pedemonte et al., 1998; Tingley and Buzsáki, 2018; Wirtshafter and Wilson, 2019). It is possible that the lateral septum also contributes to the generation of the hippocampal theta rhythm (Monmaur et al., 1993; Pedemonte et al., 1998; van der Veldt et al., 2021) in addition to the strong theta drive originating from the medial septum. The spiking activity of lateral septum neurons is correlated with locomotory behavior (running speed, acceleration, movement direction) and position (Leutgeb and Mizumori, 2002; van der Veldt et al., 2021; Wirtshafter and Wilson, 2020, 2019; Zhou et al., 1999). These studies have reported place cell-like activity and a stable spatial rate code in the lateral septum, but with a lower precision than found in the hippocampus. One study, however, suggested that instead of a rate code the lateral septum employs a phase coding strategy in which the theta phase of spikes conveys spatial information (Tingley and Buzsáki, 2018). These studies, however, did not investigate theta cycle skipping dynamics of spatial representations in the lateral septum cell population.

The lateral septum is a heterogeneous structure with regional variations in connectivity, gene expression profiles and physiology (Besnard and Leroy, 2022; Rizzi-Wise and Wang, 2021). The regional specialization may underly different functional lateral septum circuits that control a variety of motivational behaviors (Besnard and Leroy, 2022), including social aggression (Leroy et al., 2018), food seeking (Carus-Cadavieco et al., 2017; Zhang et al., 2022) and reward processing (Luo et al., 2011). The prominent hippocampal inputs are organized topographically, such that cells in the dorsal Cornu Ammonis (CA1-3) and subiculum project to dorsal-posterior parts of the lateral septum, and more ventrally located hippocampal and subicular neurons project to progressively more ventral-anterior and lateral parts of the lateral septum (Besnard and Leroy, 2022; Leroy et al., 2018; Rizzi-Wise and Wang, 2021; Tsamis et al., 2020; van der Veldt et al., 2021). This organization of hippocampal input is reflected in the physiology of lateral septum neurons, with stronger spatial coding in the dorsal-posterior lateral septum compared to ventral regions (van der Veldt et al., 2021). However, whether the theta dynamics of spatial representations in the lateral septum show subregional variations is currently unknown.

In this study we investigated the temporal dynamics of spatial representations in the lateral septum. Neuropixels probes were used to record along the dorsal-ventral extent of the lateral septum while animals performed a spatial navigation task. The activity of a large fraction of cells was theta rhythmic and a subset of cells showed evidence of theta cycle skipping. Both theta rhythmicity and cycle skipping were most prominent in the dorsal lateral septum. Using neural decoding, we show that the lateral septum cell population encodes both the current location and alternatingly the possible future paths within single theta cycles. Taken together, these findings indicate that hippocampal spatial representations and their temporal dynamics are conveyed to subcortical projection areas.

## Results

### Neural recordings across different septal subregions during execution of alternation/switching task

To characterize the temporal and spatial coding properties of neurons in the lateral septum (LS), we recorded activity along the dorsal-ventral extent of the LS spanning both dorsal (LSD) and intermediate (LSI) subregions (**Figure 1A, B**; **Figure 1 – figure supplement 1**) in four rats. Animals were trained on a Y-maze (**Figure 1C**) to run between a home (H) platform and left (L)/right (R) goal arms as part of a spatial alternation task (**Figure 1D, left**) or switching task (**Figure 1D, right**). Both tasks were continuous in nature without imposed discrete trial structure. When we refer to a “trial” below, we mean a completed lap from home to goal and back home.

**Figure 1.**
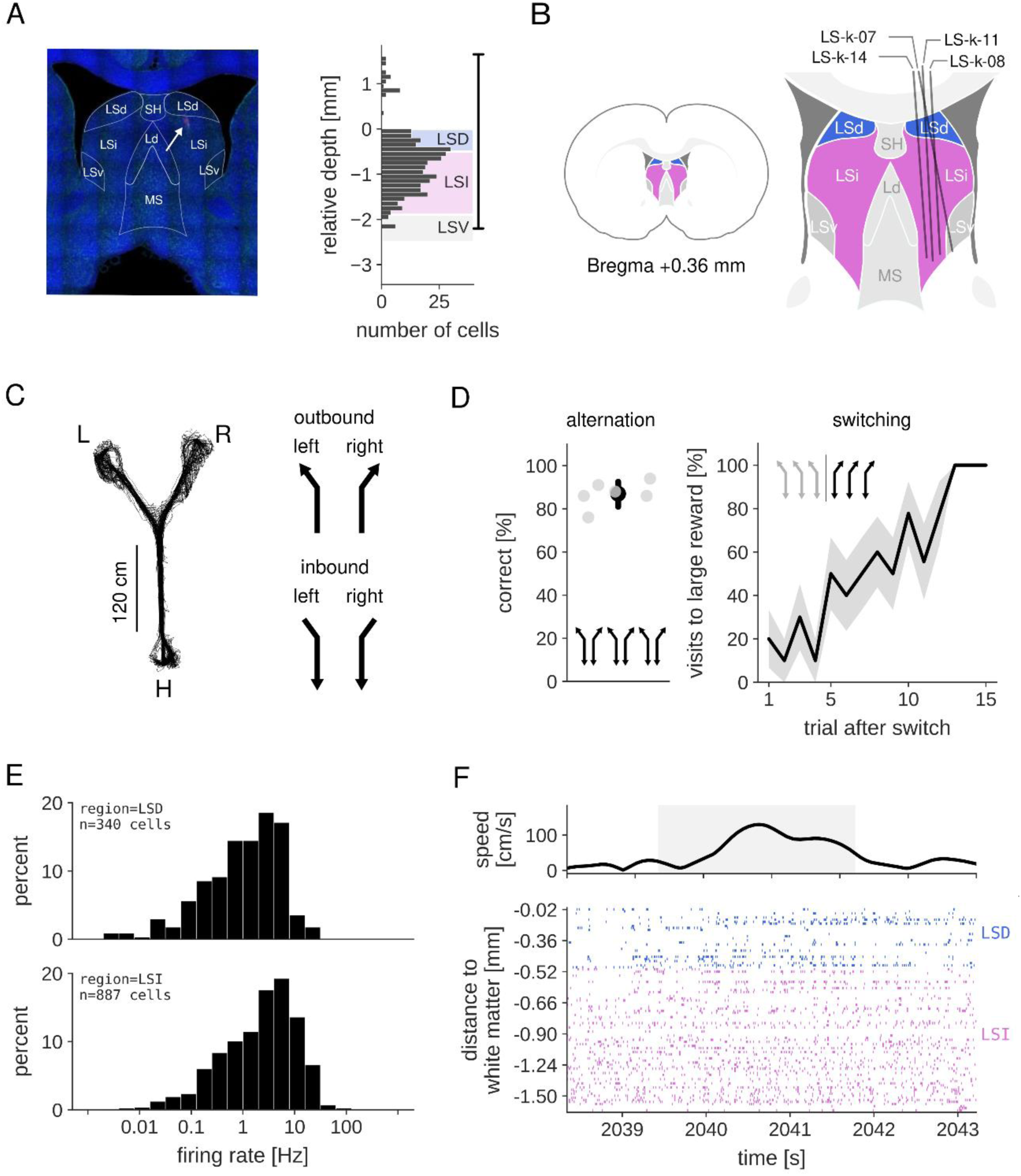
Neural recording in the lateral septum. (A) Left: coronal brain slice showing part of the Neuropixels probe trajectory (white arrow) in the lateral septum (animal LS-k-7). Right: the number of recorded cells along the probe shank for one recording session in the same animal. Depth is measured relative to the white matter above the lateral septum. Vertical line at the right indicates the span of the recorded electrodes on the probe. (B) A schematic overview of the probe tracks in all animals, projected onto a single coronal slice at 0.36 mm anterior to Bregma. (C) Left: example tracked position from a session in which the animal performed the alternation task on the Y-maze. Right: definition of outbound and inbound trajectories. (D) Left: the percentage of correct visits for all alternation task sessions. Gray dots represent individual sessions, black dot represents median and 95% confidence interval across sessions. Right: the mean percentage of correct visits for 15 trials after a reward switch for all switching task sessions. (E) Log-distribution of mean firing rates for LSD (top) and LSI (bottom) cells in all analyzed sessions. Overall mean±sem firing rate: LSD 2.52±0.18 Hz, LSI 5.21±0.24 Hz. (F) Example behavior and spiking activity in a single session during one outbound journey. Top: running speed. Gray region marks the outbound journey. Bottom: spike raster plot of all cells recorded in LSD and LSI.

In the alternation task, rats received rewards for alternatingly visiting the two goal arms on a trial-by-trial bias. In the switching task, one of the goal arms was associated with a large reward and the other goal arm with a small reward. Once animal completed eight trials to the highly rewarded goal arm, the reward contingency of the goal arms was switched, and the behavior of the animals gradually followed (**Figure 1D, right**). For most analyses, data from both tasks were grouped, unless stated otherwise.

Sessions chosen for analysis were those in which the animals were familiar with the task and performed over 75% correct choices (spatial alternation task) or at least one successful arm switch (switching task). Outbound journeys that started at the home platform and ended in one of the goal arms were analyzed separately from inbound journeys in which rats ran from the goal arms toward home (**Figure 1C, right**). We analyzed a total of 1227 cells across 12 sessions from four animals (**Table 1**). The mean firing rate of LSD cells was lower than the mean firing rate of cells located in LSI (**Figure 1E**). An example of spiking activity on one outbound journey is shown in **Figure 1F**.

**Table 1.** Overview of data. An overview of all analysed sessions from four animals performing either an alternation task or a switching task. For both tasks, the number of trials is indicated, and for the switching task the number of reward contingency switches is indicated between brackets. For both septal subregions LSD and LSI, the number of recorded cells and corresponding mean firing rate are listed.

| Animal | Task |  | Brain region LSD |  | Brain region LSI |  |
| --- | --- | --- | --- | --- | --- | --- |
|  | alternation | switching | number of cells | mean rate [Hz] | number of cells | mean rate [Hz] |
| LS-k-7 | 84 |  | 29 | 4.70 | 69 | 5.25 |
|  | 46 |  | 25 | 3.43 | 79 | 4.16 |
|  | 56 |  | 34 | 3.62 | 92 | 4.56 |
| LS-k-8 |  | 40 (2) | 22 | 1.90 | 87 | 5.61 |
|  |  | 47 (1) | 32 | 2.40 | 88 | 4.53 |
|  | 39 |  | 29 | 1.55 | 72 | 5.25 |
| LS-k-11 |  | 48 (1) | 32 | 1.21 | 60 | 8.45 |
|  |  | 44 (2) | 14 | 1.16 | 62 | 4.17 |
|  |  | 56 (3) | 31 | 1.93 | 47 | 6.53 |
| LS-k-14 | 52 |  | 30 | 1.94 | 56 | 5.14 |
|  | 49 |  | 32 | 2.88 | 78 | 5.23 |
|  |  | 51 (3) | 29 | 2.84 | 93 | 4.96 |
| Total | N=6 sessions | N=6 sessions | N=340 cells | 2.52 | N=887 cells | 5.21 |

### Strong theta rhythmicity in the lateral septum

Lateral septal neurons receive rhythmic theta frequency (6-10 Hz) input from the hippocampus and their firing is locked to theta oscillations in the local field potential (Bender et al., 2015; Tingley and Buzsáki, 2018; Wirtshafter and Wilson, 2019). We first characterized the theta rhythmic firing pattern of lateral septum neurons during outbound and inbound journeys in our dataset. Theta rhythmicity was present in the spiking auto-correlogram of cells in both LSD and LSI subregions (**Figure 2A**). To quantify the level of theta rhythmicity, we computed the relative theta peak power in the spectrum of binned spike trains (**Figure 2B**) and classified cells as theta rhythmic if the relative peak power was significantly higher than the value expected from locally jittered spike trains (see Methods for details; **Figure 2B**). More than half of the cells in LSD and LSI showed significant theta rhythmic firing (LSD: 56.9%, LSI: 59.2%; see **Figure 2 – figure supplement 1A**). The firing rate of theta-rhythmic cells was significantly higher than the firing rate of non-rhythmic cells for both subregions (**Figure 2 – figure supplement 1B**). The strength of theta rhythmicity varied along the dorsal-ventral axis of LS, such that cells located more dorsally had higher relative theta peak power than cells located ventrally (**Figure 2 – figure supplement 1C**). As a consequence, the relative theta peak power was significantly higher for cells in LSD as compared to cells in LSI (**Figure 2 – figure supplement 1D**).

### Theta cycle skipping in lateral septal cells is trajectory specific

At binary choice points in the environment, the activity of hippocampal place cell ensembles upstream of the LS switches between two alternative future paths in phase with the local theta rhythm (Kay et al., 2020). At the single cell level, this switching is evident from cells being activated at alternating theta cycles and increased rhythmicity at half theta frequency (i.e., “theta cycle skipping”). Here we asked if neurons in LS display similar cycle-to-cycle dynamics that is consistent with theta cycle skipping behavior.

In a subset of LSD and LSI cells we found evidence of enhanced rhythmicity at half theta frequency (i.e., time lag of ∼250 ms) in the spiking auto-correlogram (**Figure 2A**) that corresponds to a peak in the spectrum around 4 Hz (**Figure 2B**). For each cell, we computed a cycle skipping index (CSI) as the normalized amplitude difference between the first and second theta-related peaks in the spiking auto-correlogram (Brandon et al., 2013; Kay et al., 2020). Given variability in the theta frequency over time, the auto-correlogram of most cells takes the form of a dampened oscillation and the expected CSI value for a purely theta-rhythmic (non-skipping) cell is slightly negative. CSI values were compared to a shuffle distribution that was constructed from locally jittered spike trains that maintained theta rhythmicity but destroyed cycle skipping (**Figure 3A**; see Methods for details). CSI values were considered statistically significant if the Monte-Carlo p-value obtained from the shuffle procedure was smaller than 0.05 (**Figure 3B**).

**Figure 3.**
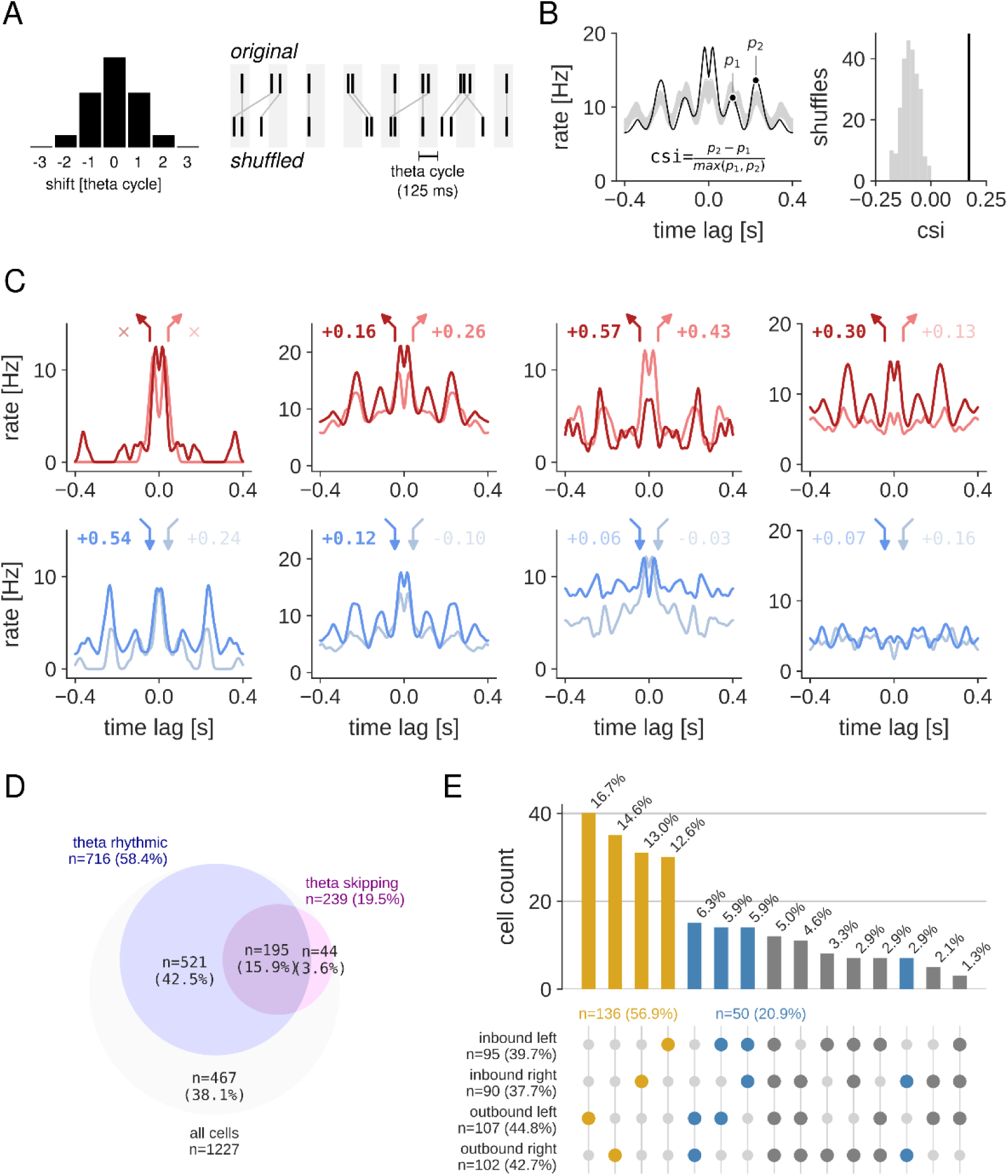
Theta cycle skipping is trajectory specific. (A) Local shuffling procedure to compute significance of cycle skipping effect in single cells. Each spike is randomly shifted by a multiple of the theta cycle (fixed to 125 ms) according to a normal distribution (shown on the left). (B) Left: example spiking auto-correlogram of original spike train (black line) and shuffled spike trains (gray). A cycle skipping index (CSI) is computed as the normalized difference between the first and second theta-related peaks. Right: distribution of CSI values for 250 shuffled spike trains (gray) and CSI value of the original spike train (black line). (C) Spiking auto-correlograms of four example cells in LSD and LSI separately for each of the four kinds of journeys (top: outbound journeys, bottom: inbound journeys). Legend at the top of each plot indicates for each journey type the CSI value (printed in bold for significant CSI values). The symbol × means that a cycle skipping index could not be computed because of low number of spikes. From left to right, the first and fourth cell show significant cycle skipping only for a single trajectory type (respectively, inbound left and outbound left). The second cell shows significant cycle skipping for three trajectory types, and the third cell for both outbound trajectories. (D) Venn diagram showing overlap of cell populations with significant theta rhythmicity and theta cycle skipping. (E) Histogram of the number of cells with a significant cycle skipping index for all possible journey combinations. Note that for most cells (56.9%; yellow) cycle skipping occurs only on a single journey type. For another population of cells (20.9%; blue), cycle skipping occurs on outbound, inbound, left, or right journeys.

We first focused on the properties of theta cycle skipping cells and noticed that in the spiking auto-correlogram the enhanced rhythmicity at half theta frequency was dependent on the type of trajectory (**Figure 3C**). In our further analysis, we divided journeys into four different trajectory categories: inbound left, inbound right, outbound left, and outbound right, and quantified theta cycle skipping only for cells that had at least 50 spikes for one or more trajectory types. For each cell, a Holm-Sidak p-value correction was applied to correct for the number of trajectories tested. We found that 27.8% (75/270) of cells in LSD and 21.5% (164/763) of cells in LSI had significantly higher CSI values than expected by chance on at least one of the trajectories (see **Figure 3 – figure supplement 1A**). As expected, the population of cycle skipping cells overlapped, to a large extent, with the population of theta rhythmic cells (**Figure 3D**). Cells located more dorsally in the LS on average showed higher CSI values than cells located in the ventral part (**Figure 3 – figure supplement 1B**). Consequently, the mean CSI values were significantly higher for cells in LSD, compared to LSI (**Figure 3 – figure supplement 1C**).

Next, we looked more closely at the selectivity of theta cycle skipping based on the type of trajectory. We noticed for most of the cells (56.9%, 136/239), the theta cycle skipping behavior happens on only one type of trajectory (**Figure 3E**). For another population of cells (20.9%, 50/239) the skipping behavior occurs on a combination of two trajectories: outbound, inbound, left, or right. This type of trajectory selectivity was consistent across both septal subregions (**Figure 3 – figure supplement 2**).

Given the trajectory dependence of cycle skipping activity, we next asked if cycle skipping occurred at specific locations on the track. To quantify the spatial preference for theta cycle skipping in individual cells, auto-correlations and CSI values were computed for spiking activity on multiple overlapping 60 cm-long subsections of the track along each of the trajectories (**Figure 4**). This analysis was only carried out for cell-trajectory pairs for which significant cycle skipping was already established using all spikes along the full trajectory.

**Figure 4.**
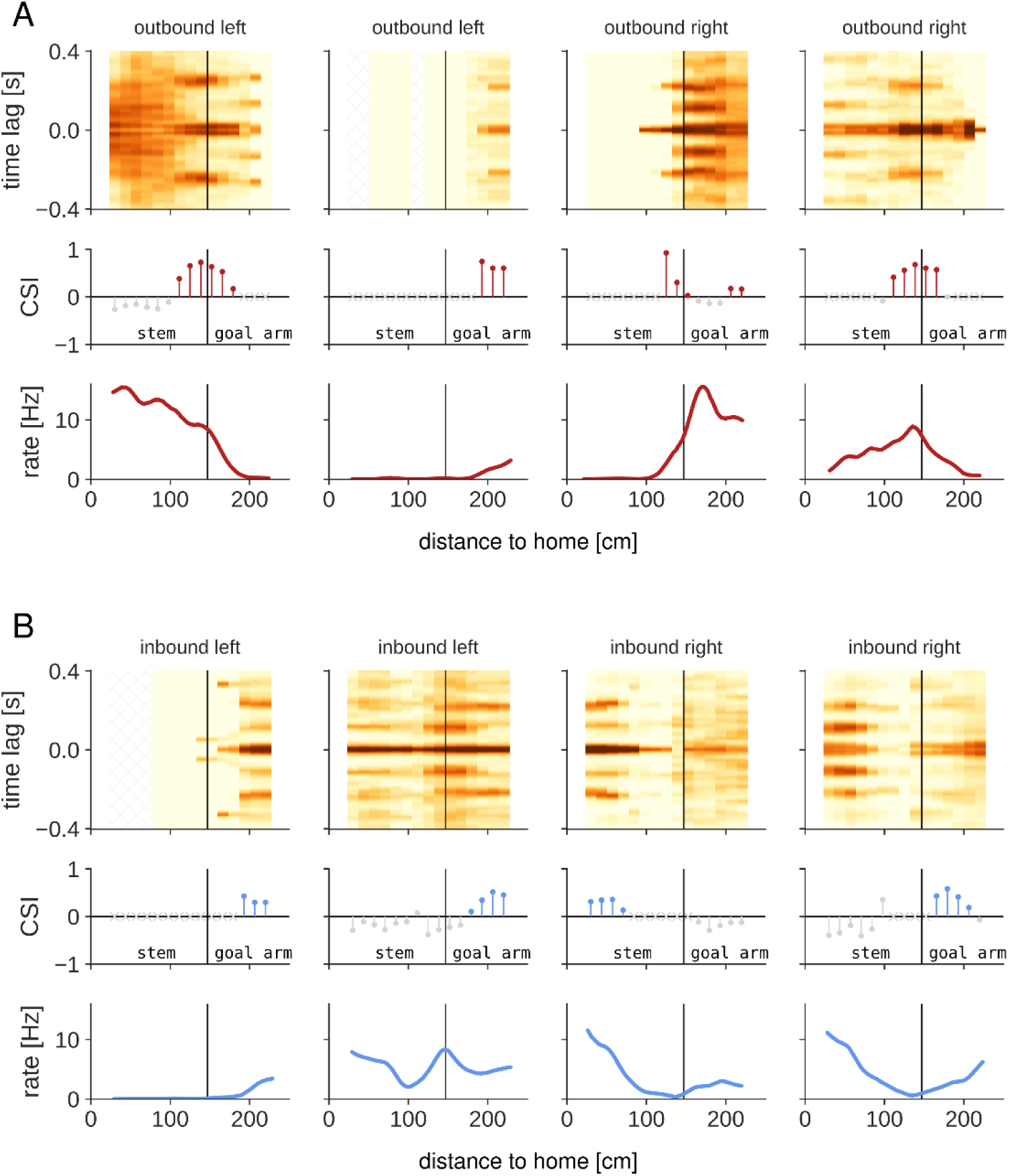
(A) Spatially resolved auto-correlation (top), corresponding cycle skipping index (CSI; middle), and spatial tuning (bottom) for four example cells during <u>outbound</u> journeys. Auto-correlation and cycle skipping index were computed for overlapping 60 cm long sections along the trajectory to the goal. In the middle plot, colored points indicate significant cycle skipping (p<0.05), light grey crosses indicate that too few spikes (<50) were available and no cycle skipping index was computed. Black vertical lines indicate the choice point separating the stem and goal arms. The first and third example were recorded in the alternation task, the other two examples in the switching task. (B) Same as (A) but for cells with significant cycle skipping on <u>inbound</u> journeys. The first and second example were recorded in the switching task, the other two examples in the alternation task.

Individual cells showed cycle skipping mainly around the choice point and in the goal arm on outbound and inbound trajectories (**Figure 4**). Across all cell-trajectory pairs, CSI values on outbound journeys were high right before the choice point and in the goal arm (**Figure 5A,B left**). On inbound journeys, CSI values were high in the goal arm leading up to the choice point (**Figure 5A,B right**). These higher CSI values corresponded to a higher percentage of cell-trajectory combinations with significant cycle skipping as compared to jittered spike trains (**Figure 5C**). These data show that cycle skipping occurs specifically around the choice point and in the goal arm in both run directions.

**Figure 5.**
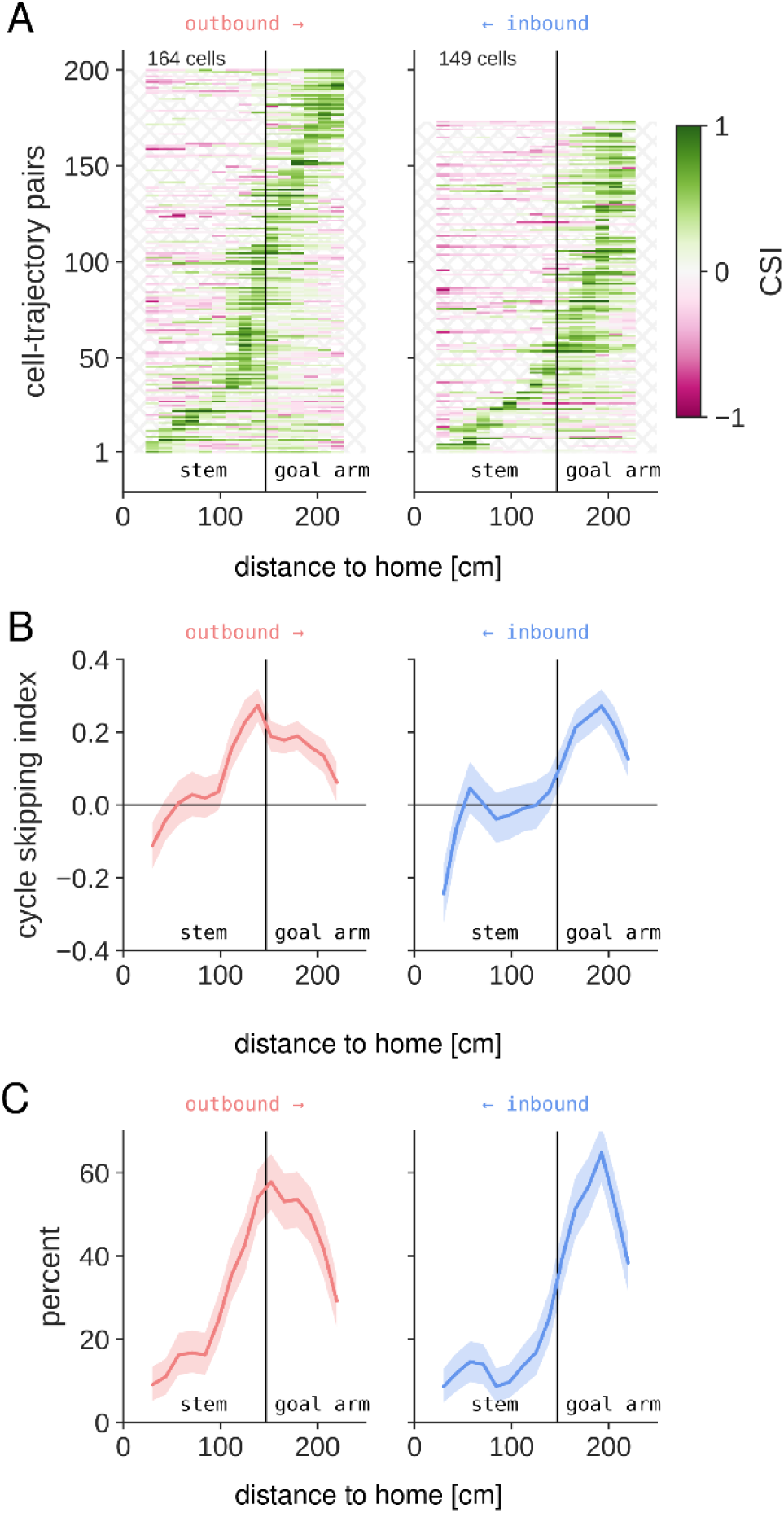
Theta cycle skipping occurs on approach to the choice point and in the goal arms. (A) Spatially-resolved (overlapping 60 cm bins) cycle skipping index for all outbound (left) and inbound (right) cycle skipping cells (in both tasks) with a minimum of 50 spikes in at least one of the spatial bins. Left/right trajectories were analyzed separately. Spatial bins that do not meet the minimum number of spike requirement are not shown, exposing the background hatching pattern. Cell-trajectory pairs are sorted according to the location of their maximum CSI value. (B) Average CSI value as a function of location along outbound (left) and inbound (right) trajectories. Thick lines represent average across all analyzed cells-trajectory pairs to/from left and right goals. The shaded region represents 95% CI. (C) Same as (B) but for the percent of analyzed cells-trajectory pairs with significant cycle skipping.

Theta cycle skipping cells showed diverse spatial firing characteristics (see also next section on spatial coding) and the locations with high CSI values did not necessarily correspond to the highest firing rate (e.g., see 1^st^ example in **Figure 4A**). To test if theta cycle skipping cells were active preferentially near the choice point or other locations, we computed the mean-corrected spatial tuning curves for cell-trajectory pairs with and without significant theta cycle skipping. We observed that, on average, the population of theta cycle skipping cells showed higher firing rate in the goal arms than in the stem of the maze as compared to non-skipping cells for outbound and inbound directions (**Figure 5 – figure supplement 1**).

### Spatial coding in the lateral septum

Given that theta cycle skipping cells in lateral septum are highly trajectory specific, we next investigated spatial properties of lateral septal cells and the relation to cycle skipping dynamics. In previous studies it was shown that lateral septal neurons convey spatial information (Leutgeb and Mizumori, 2002; Takamura et al., 2006; Wirtshafter and Wilson, 2020, 2019), although less than hippocampal neurons. Consistent with these studies, the firing rate of a large fraction of LS cells in our dataset was non-uniform across position in the Y-maze with spatial information higher than expected by chance (**Figure 6 – figure supplement 1**; percent spatially modulated cells, LSD: 75.0%, LSI: 59.4%). Even so, spatial information was generally low (<1 bit/spike), indicating only modest spatial tuning that was stronger for cells in LSD as compared to LSI (**Figure 6 – figure supplement 1**).

Spatial tuning curves for each neuron were computed separately for the four different trajectories (i.e., outbound left/right and inbound left/right; **Figure 6A**). We observed that a subset of septal neurons showed trajectory-specific activity, with cells firing differentially on the left/right goal arms (**Figure 6A, top row**) or in the outbound versus inbound journeys **Figure 6A, bottom row**).

**Figure 6.**
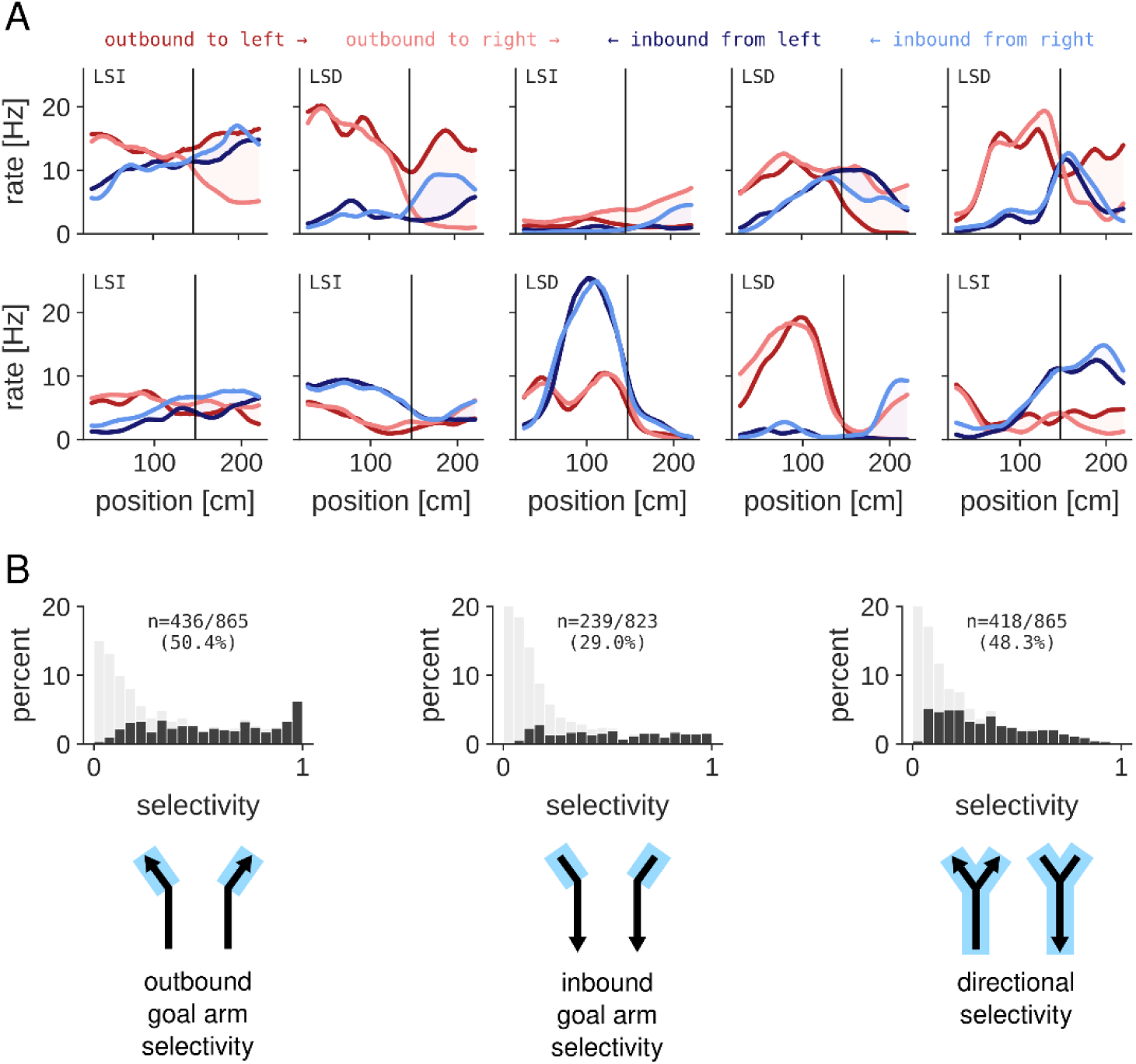
Spatial coding in the lateral septum. (A) Spatial tuning curves for 10 example neurons in LSD and LSI for the four different trajectories. (top: trajectory-specific neurons firing differentially on the left/right goal arms, bottom: direction-specific neurons firing differentially on the outbound versus inbound journeys). Examples in top row and right most example in bottom row are recorded in the alternation task, other examples are recorded in the switching task. (B) Distribution of outbound goal arm selectivity (left), inbound goal arm selectivity (middle), and directional selectivity (right) of all analyzed neurons in LSD and LSI. Light gray: full distribution of all cells. Black: highlighted part of the distribution that represent cells with significant selectivity (Monte-Carlo p-value < 0.01).

We quantified the goal arm selectivity for outbound and inbound journeys separately for active cells with a mean firing rate higher than 1 Hz on at least one of the goal arms. Overall, a high percentage of all cells were active on one or both goal arms (percentage of cells, outbound: 70.5% (865/1227), inbound: 67.1% (823/1227)). Of the active cells, the fraction that showed a significantly higher firing rate on one of the goal arms was higher in the outbound direction as compared to the inbound direction (**Figure 6B**; outbound: 50.4% (436/865), inbound: 29.3% (239/823)).

When split by subregion, a higher percentage of cells in LSI were active on the goal arms than cells in LSD (percentage of cells, outbound: LSD 60.6% (206/340), LSI 74.3% (659/887), inbound: LSD 57.4% (195/340), LSI 70.8% (628/887)). However, of the population of goal arm active cells, a higher fraction of cells in LSD than in LSI was selective for one of the goal arms (i.e., cells fired at a higher rate on one of the goal arms), and higher in outbound journeys as compared to inbound journeys for both brain regions (**Figure 6 – figure supplement 2A,D**). Within the population of goal arm selective cells, the relative spiking rate difference between the left and right arms was higher for cells located more dorsally in the LS for both outbound and inbound journeys (**Figure 6 – figure supplement 2B,E)**. This gradient of goal arm selectivity resulted in a significant higher selectivity for goal-arm selective cells in LSD than in LSI (**Figure 6 – figure supplement 2C,F**).

In contrast to the strong goal-arm specific activity, lateral septal neurons showed little prospective or retrospective coding on the stem of the Y-maze (**Figure 6 – figure supplement 3A,B**). We tested the ability to decode the outbound trajectory type (i.e., left or right outbound) from the population activity in the stem and found a classification performance below 60% (**Figure 6 – figure supplement 3C**), consistent with limited prospective coding. For the inbound trajectories, classification of the left/right trajectory in the stem was variable and on average around 60% (**Figure 6 – figure supplement 3D**). In contrast, classification performance of run direction (outbound/inbound) was high in both the stem and goal arms (**Figure 6 – figure supplement 3E**).

We next analyzed the relative firing rate difference in outbound and inbound trajectories (i.e., directionality) of LS neurons. Of the cells that were active with a mean rate above 1 Hz in either outbound or inbound direction (percentage of cells, 70.5% (865/1227)), approximately half showed directional tuning (**Figure 6B**; percentage of cells, 48.3% (418/865)). Similar to goal arm selectivity, we found more direction-selective cells in LSD as compared to LSI (**Figure 6 – figure supplement 2G**). Within the population of direction-selective cells, the strength of directionality depended on the cell location along the dorsal-ventral axis (**Figure 6 – figure supplement 2H**). Consequently, the mean directionality was significantly higher in LSD than LSI (**Figure 6 – figure supplement 2I**). The directional and goal arm selective neurons formed discrete but partially overlapping cell populations (**Figure 6 – figure supplement 2J,K**).

### Stronger position and direction coding in the dorsal lateral septum cell population

Given the spatial coding properties of individual LSD and LSI cells, we tested how well both position and running direction were represented in the population of cells. For this, we performed neural decoding analysis on outbound and inbound journeys and quantified the decoding error (**Figure 7**). Overall, there was a good correspondence between the estimated and true position and running direction when using all LSD and LSI cells for decoding (**Figure 7A,B**). The median error for position ranged from 6.0–18.0 cm across sessions (mean of all sessions: 10.2 cm; **Figure 7C**) and the percentage correct for running direction ranged from 86.9%-96.2% (mean of all sessions: 91.5%; **Figure 7C**).

**Figure 7.**
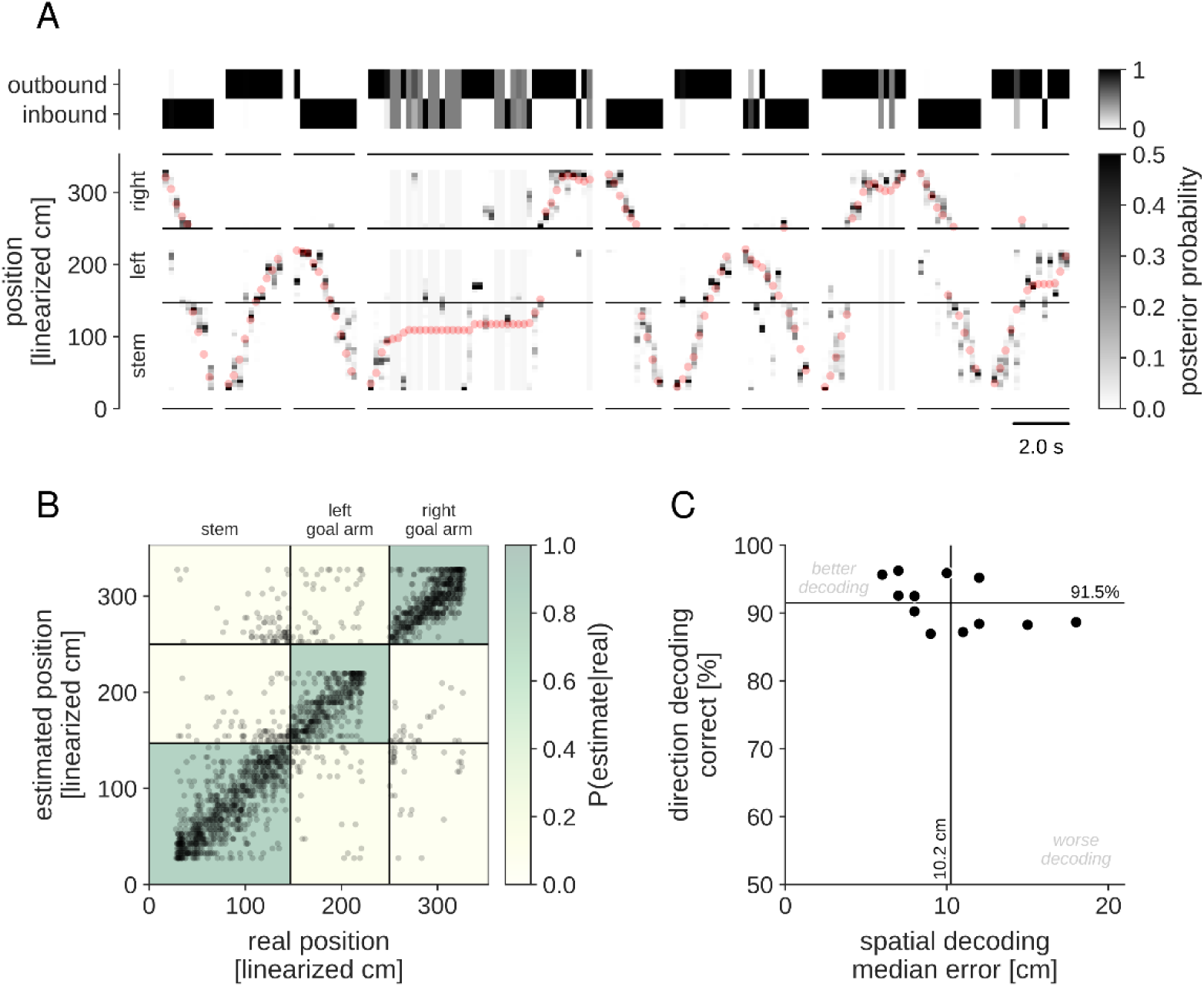
Position and direction coding in the lateral septum cell population. (A) Example result of decoding run direction (top) and position (bottom) for a single dataset (animal LS-k-7). For both direction and position, the marginal posterior probability is shown in grey scale. Position on the track is “linearized” and the horizontal black lines indicate the extent of the three maze sections: stem (bottom), left goal arm (middle) and right goal arm (top). Note that the home platform and goal platforms are excluded from the encoding model, and no decoding is performed for the time that the animal spent at the platforms. A sequence of ten outbound/inbound journeys is shown (separated by vertical lines). Red dots indicate the true position of the animal on the track. (B) Confusion matrix of the decoding result for the same session as in (A). Each dot represents a single maximum a posteriori position estimate in a 200 ms time bin during run periods (speed > 10 cm/s) in outbound/inbound journeys. The diagonal structure indicates good correspondence between estimated and true positions. The color map in the background shows the confusion matrix for decoding the three maze sections. For this session, median position error is 12.0 cm and 94.7% of direction estimates are correct. (C) Spatial and direction decoding performance for all sessions using all LSD and LSI cells combined. Vertical and horizontal lines indicate mean performance across sessions.

We next looked at position and direction coding separately for the populations of LSD and LSI cells. Using all available cells, decoding performance was similar for the LSD and LSI cell populations. However, the number of LSD cells in our dataset is approximately half of the number of LSI cells. When decoding performance was characterized as a function of the number of cells, fewer LSD cells than LSI cells were required for a similar performance level (**Figure 8A,B**). To quantify the difference in decoding performance between the two regions, we sampled a fixed number of cells for each session (n=20 cells). Both the direction and spatial decoding performance was significantly higher for LSD than LSI (**Figure 8C**).

**Figure 8.**
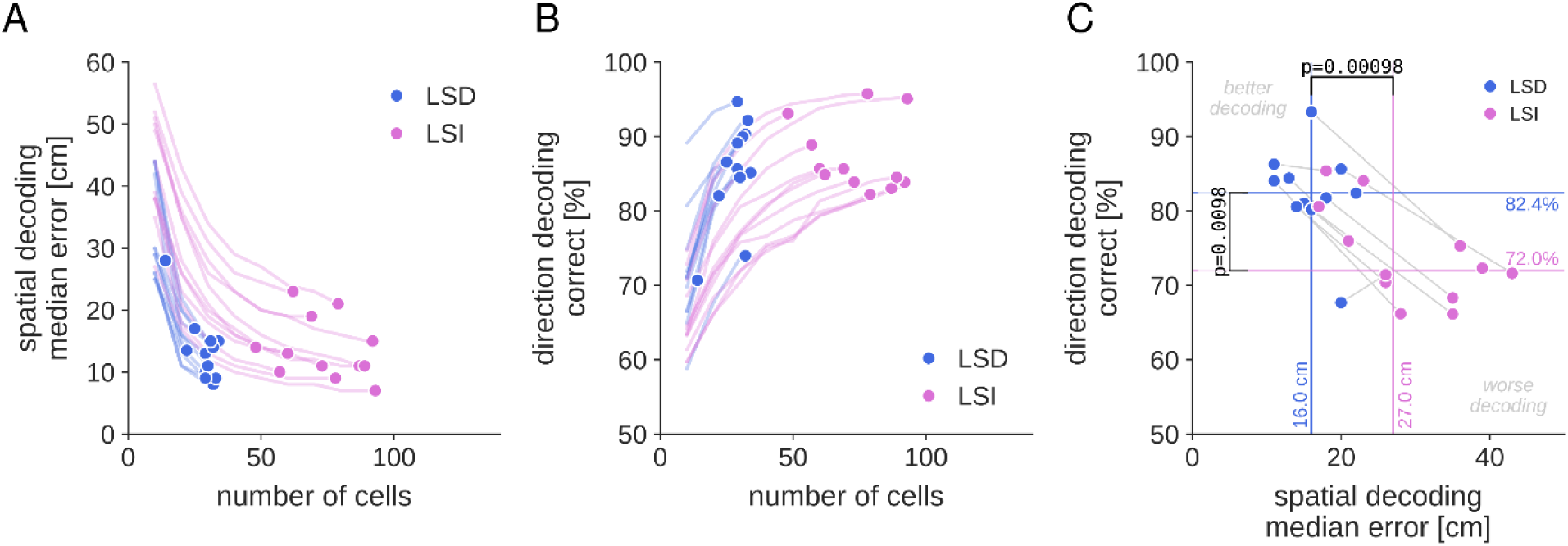
Spatial coding is stronger is LSD than LSI. (A,B) Dependency of decoding performance for position (A) and direction (B) on the number of cells included in the decoding model. The result for each session is plotted separately. Dots: the decoding performance for each session when including all available cells. Lines: the decoding performance for each session when subsampling the cells. Note that for a similar decoding performance, fewer LSD cells than LSI cells are required. (C) Comparison of decoding performance between LSD and LSI for a fixed cell population (n=20 cells). Each connected pair of dots represents a single session. Horizontal and vertical lines represent the population mean. Wilcoxon signed rank test, position: statistic=0.0, p=0.00098; direction: statistic=5.0, p=0.0098.

### Theta-cycle dynamics of lateral septum spatial representations

We next analyzed the decoded spatial representations in the population of lateral septum cells at the timescale of theta cycles. For this, we used the same encoding model as before, but decoding was performed in overlapping 50 ms time bins. As expected, spatial and directional estimates are noisier at this finer timescale, but still largely reflect the behavior of the animal (**Figure 9**). In addition, we observed non-local spatial representations of the two maze arms where the animal was not located, most strongly when animals ran towards the choice point (outbound on the stem, or inbound on the goal arms; see **Figure 9 – figure supplement 1**). To analyze the temporal dynamics of the local and non-local spatial representations, we first computed the time course of the total posterior probability *P_arm_(t)* assigned to each of the three maze arms (**Figure 9, bottom**). Next, auto– and cross-correlations of these signals were computed separately for journeys towards the choice point in either the stem (**Figure 10A**) or left/right goal arms (**Figure 10B**).

**Figure 9.**
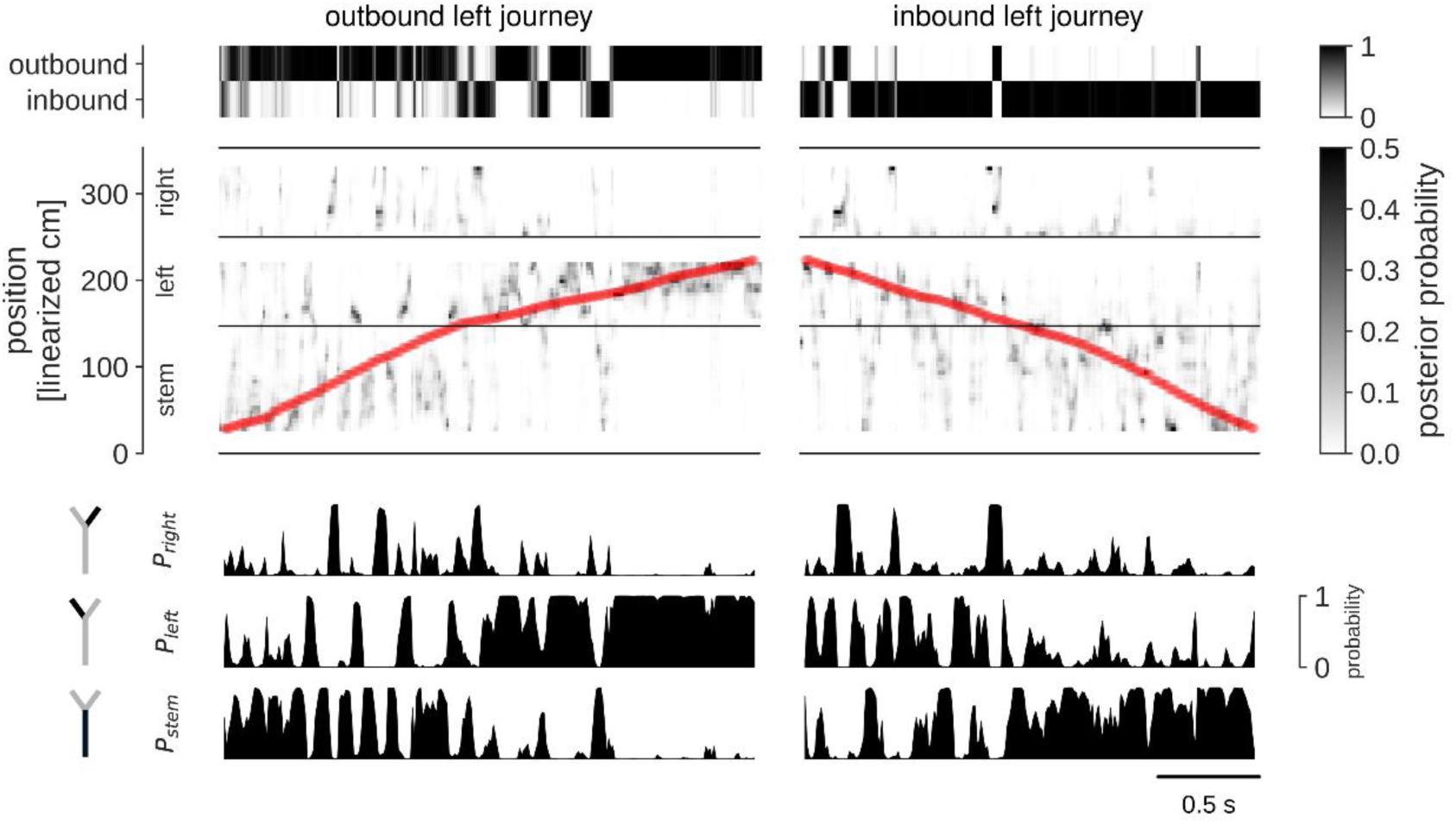
Neural decoding of theta-scale dynamics in lateral septum. Example result of decoding run direction (outbound or inbound; top) and position (middle) at fine time scale (20 ms) for a single dataset (animal LS-k-7). For both direction and position, the marginal posterior probability is shown in grey scale (black represents probability of 1 for direction, and ≥0.5 for position). Position on the track is “linearized” and the horizontal black lines indicate the extent of the three maze sections: stem, left goal arm and right goal arm. Note that the home platform and goal platforms are excluded from the encoding model, and no decoding is performed for the time that the animal spent at the platforms. A sequence of one outbound (left) and one inbound journey (right) is shown. Red dots indicate the true position of the animal on the track. Bottom: time courses of the summed posterior probability in the three maze sections.

**Figure 10.**
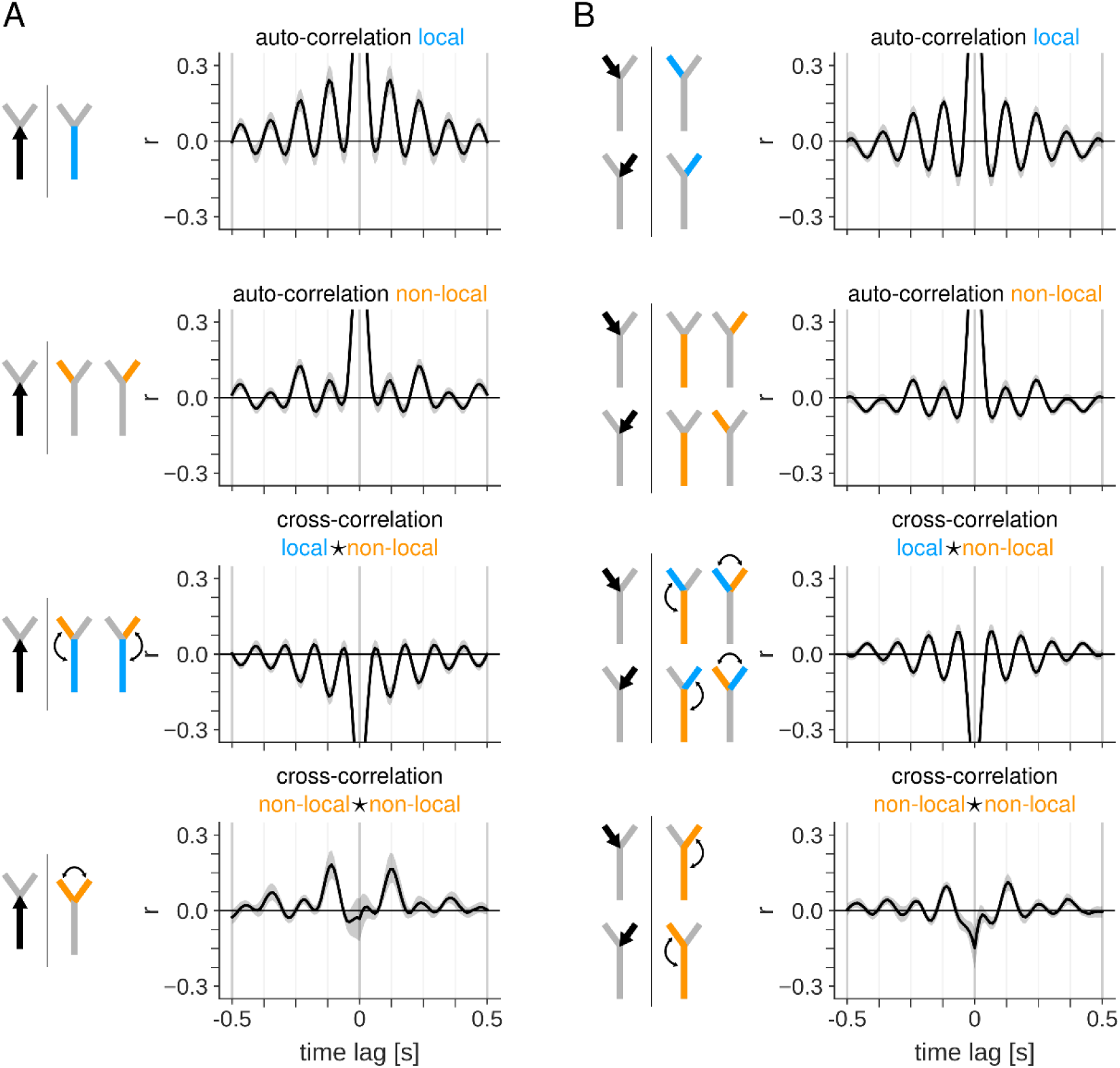
Alternation of spatial representations on approach to choice point. (A) Auto– and cross-correlation of posterior probability time courses for the three maze sections when the animal is running along the stem in the outbound direction towards the choice point. Correlations are computed as the Pearson correlation coefficient at varying time lags. For each plot, drawings at the left show the animal’s behavior (black arrow) and the maze sections for which correlations are computed (color indicates whether the highlighted maze section is local (blue) or non-local (orange) relative to the animal’s position on the track). Top: auto-correlation of local representations in the stem. Second from top: auto-correlations of non-local representations in the two goal arms. Third from top: cross-correlations between local and non-local representations. Bottom: cross-correlation between non-local representations in the two goal arms. (B) Same as (A), but for times when the animal is running along one of the goal arms in the inbound direction towards the choice point. Equivalent correlations for the two goal arms are computed jointly.

As animals ran towards the choice point in either outbound or inbound directions, the local spatial representations were strongly theta modulated (**Figure 10A,B – autocorrelation local**). In contrast, non-local spatial representations of the goal arms showed modulation at half theta frequency (**Figure 10A,B – autocorrelation non-local**), consistent with theta cycle skipping of spiking activity at the single cell level. To compare the cycle skipping dynamics of local and non-local representations, an adapted cycle skipping index was defined that quantified the normalized difference between the maximum correlation values for the single and double theta-period time lags. The cycle skipping index was significantly higher for the non-local representations than the local representations, and not different for the two run directions (**Figure 10 – figure supplement 1A**).

Cross-correlation of the local and non-local spatial representations showed an out-of-phase theta-rhythmic relationship (**Figure 10A,B – cross-correlation local ★ non-local)**, which is consistent with a model in which half of the theta cycle is used for representing the past or current location and the other half for scanning potential future locations (Kay et al., 2020; Wang et al., 2020). Finally, cross-correlation between the two non-local spatial representations (i.e., the two goal arms for outbound trajectories, and the stem and one goal arm for inbound trajectories) showed a peak at ∼125 ms (**Figure 10A,B cross-correlation non-local ★ non-local**), consistent with an alternating representation of the two possible future locations. Comparison of the peak value between the two run directions indicated a significantly stronger alternation of non-local representations in the outbound direction (**Figure 10 – figure supplement 1B**).

No pronounced theta scale dynamics of the spatial representations were observed when animals ran away from the choice point (and towards reward platforms) on either the stem (**Figure 10 – figure supplement 2A**) or goal arms (**Figure 10 – figure supplement 2B**).

### Theta cycle skipping and alternation of spatial representations is more prominent in alternation task

We next asked if theta cycle skipping at the single cell level and theta time scale dynamics of spatial representations at the population level differed between the alternation task and the switching task. While in both tasks the animals visit left and right goal arms approximately equally during a session, the most highly rewarded goal arm changes from outbound journey to outbound journey in the alternation task, whereas it remains the same during a block of trials (until a switch) in the switching task. We split the recording sessions in two groups according to the behavioral task (n=6 for both tasks) and repeated the analysis of theta cycle skipping and temporal correlations of spatial representations.

We found that a higher fraction of cells showed evidence of theta cycle skipping in the spiking auto-correlogram during execution of the alternation task than the switching task (**Figure 11 – figure supplement 1**). The location on the maze where theta cycle skipping occurred appeared to differ between the two tasks, with more pronounced cycle skipping in the alternation task just ahead of the choice point in either direction. In particular, we observed a higher fraction of cells in the alternation task with theta cycle skipping on approach to the choice point in the outbound direction (**Figure 11A,B**).

**Figure 11.**
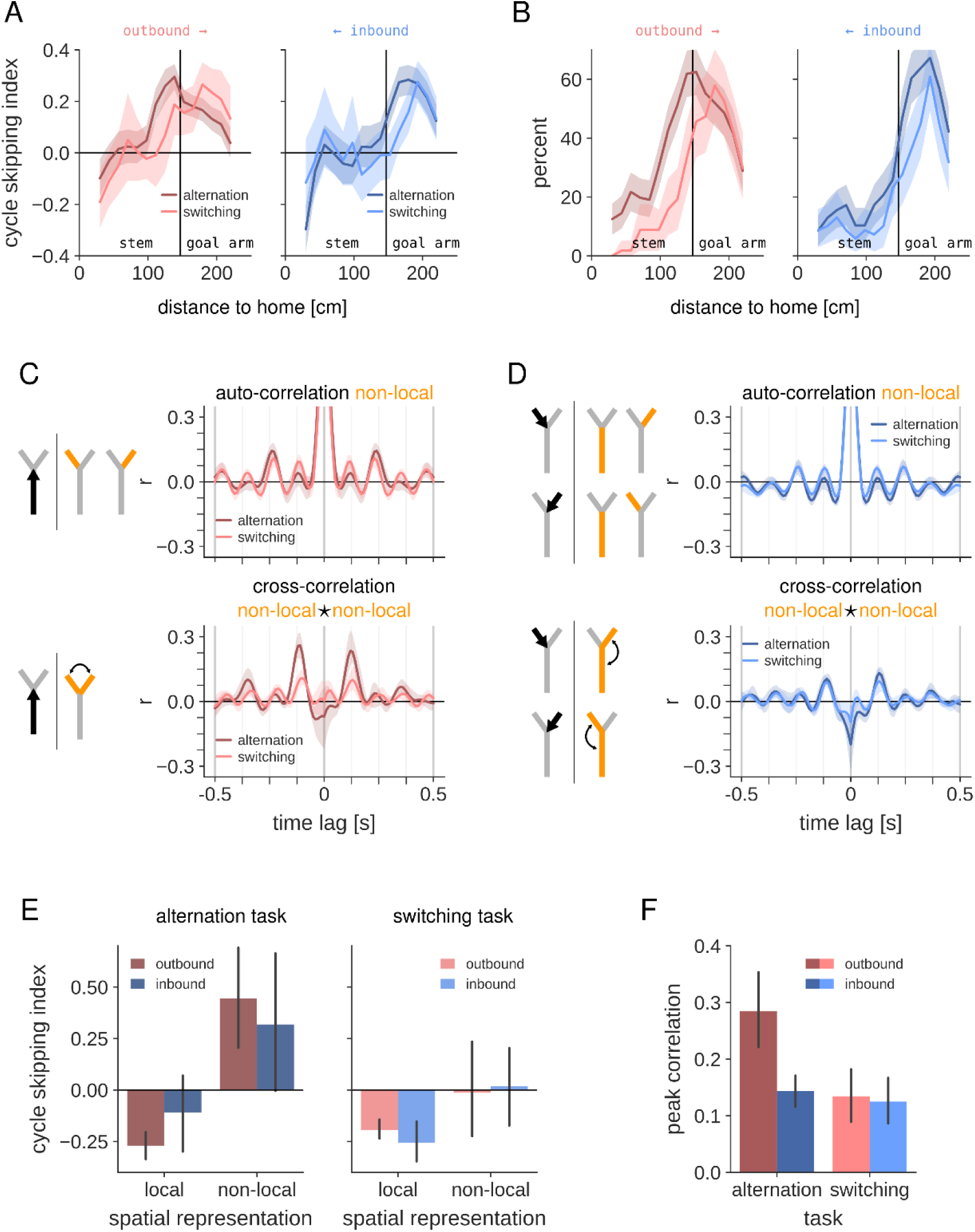
Alternation of spatial representation is task-dependent. (A) Average CSI value as a function of location along outbound (left) and inbound (right) trajectories, separately for alternation and switching tasks. Shaded region represents 95% CI. (B) Same as (A) but for the percent of analyzed cells and trajectories with significant cycle skipping. (C) Auto– and cross-correlations of non-local spatial representations in the two goal arms as animals run outbound in the stem towards the choice point, computed separately for alternation and switching tasks. See Figure 10 for details. (D) Same as (C), but for times when the animal is running along one of the goal arms in the inbound direction towards the choice point. (E) Quantification of the cycle skipping index computed from the auto-correlation of the local and non-local spatial representations for both run directions and separated by task. The results of a three-way ANOVA to evaluate the effects of task, locality, and run direction on the cycle skipping index are presented in Figure 11 – table supplement 1. (F) Quantification of the peak cross-correlation of the non-local spatial representations for both run-directions and separated by task. The results of a two-way ANOVA for the effects of run direction and task on peak correlation are presented in Figure 11 – table supplement 3. Corresponding post-hoc comparisons are listed in Figure 11 – table supplement 4.

We re-analyzed the temporal dynamics of decoded spatial representations as animals approached the choice point to compare the two behavioral tasks (**Figure 11C,D auto-correlation non-local**). The cycle skipping in the auto-correlation of the non-local representations appeared more pronounced in the alternation task than in the switching task. A three-way ANOVA was performed to evaluate the effects of task, locality, and run direction on the cycle skipping index (**Figure 11 – table supplement 1**). The results indicated a significant interaction between locality and task, in addition to significant main effects for locality and task. Post-hoc analysis of the locality ✻ task interaction shows that the cycle skipping index is significantly higher for the non-local representations in the alternation task than the non-local representations in the switching task (**Figure 11 – table supplement 2**). In addition, whereas for the alternation task the cycle skipping of the non-local representation is significantly stronger than the local representations, this was not the case for the switching task (**Figure 11 – table supplement 2**).

Alternation of the two non-local representations – as evidenced by a cross-correlation peak at theta time lag (**Figure 11C,D cross-correlation non-local ★ non-local**) – was stronger in the alternation task than switching task. We performed a two-way ANOVA for the effects of run direction and task on the cross-correlation peak value, and found a significant interaction between run direction and task, as well as significant main effects for run direction and task (**Figure 11 – table supplement 3**). Post-hoc comparisons between all combinations of groups revealed that the cross-correlation peak in the outbound direction of the alternation task was significantly larger than the peak for the inbound direction, and significantly larger than the peaks for either direction in the switching task (**Figure 11 – table supplement 4**).

These results indicate that the theta-cycle skipping dynamics of non-local spatial representations are more pronounced when animals performed the spatial alternation task. A possible explanation for the difference between the two tasks is that only the future paths that are relevant for the animal are preferentially represented in alternating theta cycles. Thus, in the switching task, as the animals develop a preference for the goal arm with the large reward, the goal arm with the small reward becomes less relevant. If this explanation is correct, than one prediction is that enhanced theta-cycle skipping dynamics will be observed in the switching task during periods in which the rats explore both goal arms before developing a bias to one goal arm. To explore this prediction, we analyzed the theta cycle dynamics of non-local spatial representations in the first trials of each session, and in the trials around the change in reward contingency in the switching task. We restricted our analysis to the outbound approach to the choice point, where we observed the strongest theta cycle skipping and alternation of non-local representations.

Contrary to the prediction, the auto– and cross-correlations of the non-local spatial representations in the first five trials of switching task sessions showed no evidence of theta cycle skipping or theta cycle alternation, in contrast to the first five trials in the alternation task (**Figure 11 – figure supplement 2**). Similarly, quantification of the theta cycle skipping and alternation dynamics in the five trials following the reward contingency change showed no significant difference with the five trials preceding the reward change (**Figure 11 – figure supplement 3**). If anything, a trend towards stronger theta cycle skipping/alternation in the trials before a reward change was observed.

Overall, these results indicate that the difference that was observed between the two behavioral tasks is not explained by the number of relevant paths available to the animal.

## Discussion

Our findings reveal that the lateral septum contains a population of neurons that alternatingly represents upcoming locations during adjacent theta cycles when rats approach a choice point. At the single cell level, this is supported by theta rhythmic spiking activity that is modulated at half theta frequency (i.e., theta cycle skipping). Our data shows that the alternating expression of spatial representations in the lateral septum is task dependent, which suggests that task demands and experience shape which representations are activated near a choice point. The lateral septum receives strong input from hippocampal place cells (Risold and Swanson, 1997), and while there may be integration and transformation of incoming spatial signals (Tingley and Buzsáki, 2018), the theta-cycle dynamics of spatial representations appear to be transferred intact.

Our findings indicate that cells in the LSD region exhibit a significantly greater degree of theta cycle skipping than cells in the more ventrally located LSI region. This regional difference in skipping is accompanied by a lower firing rate, stronger theta rhythmicity, and more pronounced spatial coding in the LSD region (conform van der Veldt et al., 2021). The most likely source of the differences between LSD and LSI is the input they receive from the hippocampus. Whereas the dorsal hippocampal CA1 and CA3 regions project to the LSD, the LSI receives input from more ventrally-located hippocampal regions (Risold and Swanson, 1997; Rizzi-Wise and Wang, 2021). Theta dynamics vary along the dorsal-ventral axis of the hippocampus, with both the power of theta oscillations and the percentage of theta-rhythmic neurons being higher in the dorsal hippocampus (Royer et al., 2010; Schmidt et al., 2013). Additionally, hippocampal spatial representations increase in scale from dorsal to ventral (Jung et al., 1994; Kjelstrup et al., 2008), and ventral hippocampal cell activity shows less differentiation between multiple arms in a maze (Royer et al., 2010). These data suggest that the differences observed between LSD and LSI are mostly derived from the upstream hippocampus. Although theta cycle skipping has not been studied in the ventral hippocampus, we predict based on our observations in the LS that it will be less prominent in the ventral hippocampus.

Reports on the percentage of cells in the lateral septum that convey spatial information in their firing rate vary considerably and range from 5% (Tingley and Buzsáki, 2018) to 56% (Wirtshafter and Wilson, 2019). The percentage of spatially modulated cells in our dataset sits at the upper end of this range. Some of this variation may be explained by the subregion that was sampled, given the higher spatial information conveyed by cells in LSD as compared to LSI, as discussed above. A second explanation for the high percentage of spatially modulated cells in our study could be the method used to identify these cells. We used a shuffling procedure that will identify also weakly modulated cells, whereas other studies have used an arbitrary cut-off of the spatial information metric (Wirtshafter and Wilson, 2019). Even though the activity of a large fraction of lateral septum cells may be spatially modulated, the information content per spike is limited, similar to previous studies in the lateral septum (Takamura et al., 2006; Wirtshafter and Wilson, 2020, 2019), and comparable to what has been reported for the ventral hippocampus (Jung et al., 1994). Nevertheless, the population activity of lateral septum neurons contains sufficient spatial information for neural decoding of the animal’s location and running direction on the maze with low error (conform van der Veldt et al., 2021).

One notable property of hippocampal spatial coding was nearly absent in the lateral septum cells, namely the trajectory specific firing (“splitter cells”) depending on the future path (prospective coding) or past path (retrospective coding) (Ferbinteanu and Shapiro, 2003; Frank et al., 2000; Wood et al., 2000). We observed a slightly stronger retrospective than prospective coding, which is in line with what has been reported in the hippocampus (Duvelle et al., 2023). In studies of trajectory-specific activity of hippocampal place cells, it has been noted that the fraction of cells that show splitter activity may depend on several factors, including the behavioral task and training experience (Duvelle et al., 2023). A previous study that also used a continuous alternation task in a 3-arm maze reported a relatively low fraction of splitter cells in area CA1 (Frank et al., 2000), which is consistent with the weak trajectory-specific activity in our dataset. It remains an open question, however, if the lateral septum is generally devoid of trajectory-specific coding or if it shows the same task-dependent variability as found upstream in the hippocampus.

The single cell level analysis of temporal spiking patterns showed that theta cycle skipping occurred as animals approached the choice point as well as in the middle of the goal arms. The population decoding analysis, however, only pointed to alternating spatial representations of upcoming arms ahead of the choice point. It should be noted that theta cycle skipping of single cells does not necessarily imply that two populations of neurons are activated in alternation. Likewise, cycle-by-cycle alternation of spatial representations at the population level can occur without observing theta cycle skipping in the spiking auto-correlation (e.g., when cells fire at a low rate). While the single cell and population analyses are consistent with each other as rats approach the choice point, it is currently not clear why an elevated theta cycle skipping is observed for individual cells in the goal arms. One explanation could be an increased excitatory drive coupled with a temporary spike-burst induced decrease of spiking probability, which is consistent with our observation of a general rate increase in the goal arm for theta cycle skipping cells. Alternatively, as our decoding analysis only looked at aggregate representations of the three maze arms, it is possible that alternating spatial representations are expressed at the population level but only of locations <u>within</u> a single arm.

Theta cycle skipping and alternating ensembles within adjacent theta cycles have been observed previously in the hippocampus (Kay et al., 2020), medial entorhinal cortex (Deshmukh et al., 2010), medial prefrontal cortex (Tang et al., 2021) and nucleus reuniens (Jankowski et al., 2014). Our result now extend this finding to the lateral septum and further point to the involvement of a large coordinated network. However, there are several notable similarities and differences between brain regions. In the hippocampus, representations of both future options appear to be expressed equally often ahead of the choice point (Kay et al., 2020; Tang et al., 2021). Our data indicates that the lateral septum behaves similarly, which is consistent with the strong direct projections from the hippocampus to the lateral septum. In contrast, while cells in the medial prefrontal cortex show strong theta cycle skipping, the corresponding theta spike sequences predicted the upcoming choice (Tang et al., 2021). Interestingly, theta cycle skipping has been observed in the nucleus reuniens, which links the medial prefrontal cortex and hippocampus (Griffin, 2015) and through this pathway mediates trajectory-dependent (prospective) activity in the hippocampus (Ito et al., 2015) and gates VTA dopaminergic activity (Zimmerman and Grace, 2016). However, given the differences between the theta cycle skipping properties of hippocampus and medial prefrontal cortex, the role of the nucleus reuniens needs further clarification. Finally, in the entorhinal cortex, theta cycle skipping has been preferentially linked to neurons that are tuned for head direction (Brandon et al., 2013), rather than place. The tendency for entorhinal cells to skip alternate theta cycles increased from dorsal to ventral entorhinal cortex (Brandon et al., 2013; Deshmukh et al., 2010), which is contrary to our observations in the lateral septum. The extent to which theta cycle skipping of head direction cells in the entorhinal cortex is functionally linked to theta cycle skipping in the hippocampal-lateral septum circuit is currently unknown and it is possible that they reflect distinct phenomena.

Individual theta cycles are often seen as a computational unit (Tang et al., 2021) that helps to segregate spatial experiences or choices (Gupta et al., 2012; Jezek et al., 2011; Kay et al., 2020). When rats are exposed to an instant switch from one environment to another (“teleportation”), there is an transient flickering between preformed representations of the two environments (Jezek et al., 2011), which may be explained by two competing attractor states in the network. Similarly, the possible future paths at fork in the road may be represented by continuous attractors in the hippocampal network that compete with each other in each theta cycle. The alternation of these attractors from cycle to cycle suggest that the ensemble that was expressed in one theta cycle subsequently may be temporarily suppressed, e.g., through firing rate adaptation (Chu et al., 2023) or through interference with a slower ∼4 Hz oscillation (Deshmukh et al., 2010), to favor the activation of another ensemble in the subsequent cycle. This may be seen as the system efficiently sampling the distribution of possible future trajectories using anti-correlated samples in subsequent theta cycles (Ujfalussy and Orbán, 2022). When there is reduced uncertainty about which future path to follow, e.g., in our switching task in which animals learn to follow one path to the high reward, the sampled distribution may be restricted to the one preferred path to reward (Johnson and Redish, 2007), consistent with a reduction in theta cycle skipping and alternation of future path representations in the switching task. Further exploration is needed to understand how and on what timescale the samples are restricted in an experience or task-dependent manner.

Theta sequences of hippocampal place cell activity that represent paths ahead of animals may support the deliberation about what action to take next. The first study that demonstrated how spatial representations sweep along left and right paths at a choice point (Johnson and Redish, 2007), linked the activity to the behavioral sampling of the two options through head movements (vicarious trial and error, or VTE). Our result show that the decoding of spatial representations in the lateral septum showed both local and non-local representations within the span of a single theta cycle, which is consistent with a sweep along the path ahead. Still, theta sweeps also occur in environments without choice points (Davidson et al., 2009; Foster and Wilson, 2007; Wang et al., 2020), and the theta sweeps that sample future options also occur on the approach to a choice point without pausing or overt VTE (this work; Kay et al., 2020). As such, theta sweeps more generally may constitute predictions of ongoing experience (Barron et al., 2020; Lisman and Redish, 2009) that could help to identify deviations from expectation (prediction error) that can be used to update stored representations or modify action plans (Howe and Blair, 2022). Our demonstration of theta cycle skipping indicates that these hippocampal predictions may engage a subcortical network for motivation and reward processing through the lateral septum.

Through the hippocampal-lateral septum pathway, the sampling of possible future paths could involve downstream hypothalamic and midbrain nuclei (Besnard and Leroy, 2022; Carus-Cadavieco et al., 2017; Deller et al., 1994; Staiger and Nürnberger, 1991; Wong et al., 2016), including the ventral tegmental area (Luo et al., 2011). These brain regions may integrate motivational and reward value components with the sampled trajectories that could support tracking of the consequences of forthcoming actions at short time scales. If this is indeed the case, we predict that alternating representations of reward or other motivational signatures can be found in the projection areas of the lateral septum.

Our finding that hippocampal theta cycle skipping and alternating theta sweeps propagate to the lateral septum, suggest that this activity engages a broad subcortical-cortical network. Whether theta sweeps in this network are required for route planning or are merely a byproduct of a circuit that is shaped by experience, may be addressed in future studies using circuit-based manipulations – e.g. inhibition of hippocampal-lateral septum projections (Bender et al., 2015) – and perturbations of theta oscillations (Mouchati et al., 2020; Petersen and Buzsáki, 2020; Quirk et al., 2021; Siegle and Wilson, 2014). For example, manipulating neural activity in alternating theta cycles could be used to preferentially suppress the expression of one of the future paths and may lead to altered choice behavior. In general, determining how the joint sampling of possible future trajectories and their outcomes contributes to learning processes in goal-oriented navigation and modulates decision-making will be an interesting direction for future studies.

## Materials and Methods

### Experimental model and subject details

A total of 4 male Long Evans rats (280g to 350g) were used for this study. All experiments were carried out following protocols approved by the KU Leuven animal ethics committee (P119/2015 and P090/2021) in accordance with the European Council Directive (2016/63/EU). Each animal was familiarized to the experimenter for two weeks prior to the start of behavioral training. Following surgical implantation of neural probes, animals were housed separately in individually ventilated cages (IVC) with ad libitum access to water and controlled intake of food pellets. To motivate animals to run for reward in behavioral tasks, animals were food deprived to no less than 85% of their free feeding body weight. Body weight and general health status were checked daily by researchers and animal care personnel.

### Behavioral training

Prior to surgery, animals were familiarized with a sleeping box and elevated linear track in the experimental room. On the linear track, animals were taught to shuttle back and forth between the two ends to collect reward (Choco-puffs). Next, animals were introduced to a Y-maze consisting of a 120 cm long stem and 76 cm long left/right goal arms. A reward platform was connected to the end of the stem (“home”) and both goal arms. Animals were not pre-trained on any task, but only placed on the maze to initiate familiarization and speed up training after the surgical procedure. Once implanted, the animals were trained over several weeks in alternation and switching tasks. After each experimental session, animals were transported to a sleep box, where they rested for 30 minutes.

In the continuous alternation task, rats could freely run and visit reward platforms for 30 minutes, but were only rewarded for outbound (home to goal) and inbound (goal to home) journeys that corresponded to the …→home→left→home→right→home→… alternation sequence. Thus, outbound journeys were only rewarded if the animals visited the goal arm opposite to the most recently visited goal arm, and inbound journeys were always rewarded. Following training, the animals rarely ran from one goal arm to the other without first returning home. We refer to a pair of completed outbound and inbound journeys as a “trial”.

In the switching alternation task, rats could freely run and visit the reward platforms, and were rewarded after each outbound and inbound journey. In one of the goal arms, the animals obtained a large reward (5 Choco-puffs) and in the other goal arm they obtained a small reward (1 Choco-puff). As the animals developed a preference to shuttle between home and the highly-rewarded goal arm, the reward values of the two goal arms were switched as soon as the animals visited the highly-rewarded goal arm eight times in a row. Only sessions with at least one switch were analyzed. The goal that was initially associated with a large reward was randomly selected at the beginning of each session. The duration of switching task sessions was set to 45 minutes, to account for the longer time it took for the animals to consume the large reward and to obtain a similar number of trials per session as for the alternation task.

### Surgical procedure

Animals were implanted with a single-shank Neuropixels probe containing 960 electrodes and 384 read-out channels (imec, Leuven, Belgium; https://ww.neuropixels.org; (Jun et al., 2017)). The probe was mounted in a 3D printed fixture that enabled reuse of the probe (van Daal et al., 2021). Immediately prior to implantation, the probe shank was disinfected with 70% isopropyl alcohol and dip-coated with fluorescent dye DiI (ThermoFisher Scientific, USA). Surgeries were performed using standard aseptic techniques. Briefly, rats were anesthetized with isoflurane (induction: 4%, maintenance: 1-2%) and injected subcutaneously with the analgesic Metacam (5 mg/kg body weight; Boehringer Ingelheim Vetmedica GmbH). Throughout the surgery, breathing rate, heart rate and blood oxygen level were continuously monitored, and body temperature was kept constant with a rectal temperature sensor and heating pad (PhysioSuite, Kent Scientific, USA). After shaving the scalp with clippers, rats were carefully placed in a stereotaxic frame. Next, the scalp was disinfected with isobetadine and isopropyl alcohol and an incision was made along the midline to expose the skull. The skull was then scraped clean and dried before 7-8 anchoring bone screws (Fine Science Tools, Heidelberg, Germany) were gently screwed into small pre-drilled holes. One bone screw served as electrical ground. A craniotomy was made above the lateral septum at 0.3 mm anterior and 0.2 mm lateral relative to Bregma. After removal of the dura mater, the Neuropixels probe was inserted to a maximum depth of 7 mm at a 3° angle in the coronal plane using a motorized micromanipulator (for detailed implantation coordinates in each animal, see Figure S1). A skull screw positioned above the cerebellum was connected to the probe and used as an reference electrode. The implant was secured in place using meta-bond (C&B super bond, Brildent, India) and light-curable dental cement (SDI wave A2, Bayswater, Australia). Post-operative analgesic (subcutaneous injection of Metacam, 5mg/kg body weight) was administered once daily for three consecutive days.

### Electrophysiological recordings

Electrophysiological data were collected using Neuropixels data acquisition hardware (imec, Leuven, Belgium) and SpikeGLX software (Bill Karsh, https://github.com/billkarsh/SpikeGLX). Position and head direction of the animal were tracked with an overhead video camera (50 Hz) using a Digilynx acquisition system and Cheetah software (Neuralynx, Bozeman, MT). In two animals (LS-k-7 and LS-k-8), two colored LEDs (red and blue) were mounted on the implant for tracking purposes. In the other animals, tracking was performed directly from the video images using DeepLabCut (Nath et al., 2019). Signals and video from the two acquisition systems were synchronized offline with the help of an external clock signal generated by an Arduino. The clock signal consisted of regular pulses at 1 Hz with varying pulse durations. The clock signal was acquired on an extra analog channel of the Neuropixels hardware and a digital input channel of the Digilynx hardware. Offline synchronization was performed by determining the optimal alignment of the pulse durations in the two recorded clock signals.

### Histology

Once the experiments were completed, animals were euthanized (Dolethal) and transcardially perfused with 4% formaldehyde following approved procedures. Brains were stored in 4% formaldehyde for 24h at 4°C after which they were transferred to 30% sucrose solution at 4°C until they were cut in 50 μm thick coronal slices with a cryostat (Leica, Germany). The brain slices were washed twice for 10 minutes with phosphate buffered saline (PBS), followed by staining with Neurotrace 500/525 green (1:300 for two hours, NeuroTrace™ 500/525 Green Fluorescent Nissl Stain – Solution in DMSO, Thermo Fisher Scientific, MA USA). Following two PBS washes, brain sections were additionally stained with DAPI (40, Diamidino-2-phenylindole dihydrochloride, Sigma-Aldrich, MA USA) for 30 minutes. Finally, brain slices were cover slipped with antifade medium (Vectashield, Vector, CA USA) and imaged on a confocal microscope (LSM 710, Zeiss, Germany) with a 10× objective.

### Quantification and statistical analysis

Data analysis was carried out using Python and its scientific extension modules (Harris et al., 2020; McKinney, 2010; Virtanen et al., 2020), augmented with custom Python toolboxes. Analysis of variance and corresponding post-hoc analysis were computed in JASP statistical software (version 0.18.10).

### Electrode locations in septal subregions

The location of the electrodes in the septal subregions was determined based on a combination of the probe implantation depth during surgery, the probe track in the histology and neural activity signatures, in particular the absence of neural activity in the white matter above the septal nuclei.

### Spike sorting

Spikes were extracted and classified using Kilosort 3 (Pachitariu et al., 2016) followed by manual curation using the software Phy (https://phy.readthedocs.io).

### Behavior

The behavior of the animals was separated into run trajectories that started and ended in one of the reward platforms at the end of the Y-maze arms. Run trajectories were categorized according to the arm of origin (e.g., left arm to stem or right arm) or according to the approach from/to the stem (outbound: from stem to left/right arm; inbound: from left/right arm to stem).

### Theta modulation

To quantify the theta modulation of single cells, spike trains were binned at a resolution of 1 ms and a multi-taper power spectrum was computed for selected behavioral epochs (parameters for spectrum: window size = 2 s, bandwidth = 1 Hz).

To compute a theta modulation index, the power peak in the theta band (6-10 Hz) was detected and a frequency window (± 1.5 Hz) around the peak was defined. The base theta power is defined as the integrated power under the line that connects the power density at the start and end of the frequency window. The peak theta power is then defined as the integrated power above this line. Finally, the theta modulation index is computed as 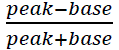. To determine if the theta modulation index was significantly higher than expected by chance, a shuffle distribution of modulation indices was computed for 500 locally time-jittered spike trains. For each spike, jitter was randomly drawn from a gaussian distribution with standard deviation of half a theta cycle (62.5 ms). A theta modulation index was considered statistically significant if the Monte-Carlo p-value was smaller than 0.01.

### Theta cycle skipping

The propensity of cells to increase their activity on alternating theta cycles was captured in a theta cycle skipping index that was calculated from the auto-correlogram (ACG) separately for each cell (Deshmukh et al., 2010; Kay et al., 2020). ACGs were computed from spikes in selected behavioral epochs, binned at 5 ms and smoothed with a gaussian window (bandwidth = 0.5 s). Cells with fewer than 50 spikes in the selected epochs were excluded from the analysis. In the ACG, the first and second theta-associated peak were detected: p_1_, the peak in the 90-200 ms window nearest to 0 ms time lag, and p_2_, the peak in the 200-400 ms window nearest to 0 ms time lag. For cases without peak p_1_ or p_2_, an ACG value was selected in lieu of the missing peak at half/double the time lag of the detected p_2_/p_1_ peak. The theta cycle skipping index was computed as 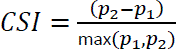.

To determine if a CSI value was higher than expected by chance, a shuffle distribution was computed from 250 locally randomized spike trains in which each spike was shifted by *k*θ, with *k* randomly drawn from *K* = {−3, −2, −1,0,1,2,3} with probability N(*K*_*i*_) and θ = 125 ms. This spike randomization procedure was designed to destroy the cycle skipping structure but keep theta modulation intact. A CSI value was considered statistically significant if the Monte-Carlo p-value was smaller than 0.05.

### Spatial tuning

The tracked animal position was converted to “linearized” location along the track by projecting the (x,y) coordinates to a skeletonized Y-maze. The position on the maze was defined as the distance from the home platform to (one of) the reward platform. For each cell, a spatial tuning curve λ(*x*) in selected behavioral epochs was then constructed by computing compressed kernel density estimates (sampled at 1 cm resolution, gaussian kernel with bandwidth 10 cm; (Sodkomkham et al., 2016)) of spike counts and occupancy as a function of the linearized position.

To quantify spatial tuning, for each cell the spatial information was computed from the spatial tuning curve according to (Skaggs et al., 1992) and either expressed in bits/spike or bits/second. To test if the spatial information measure for a given cell was higher than expected by chance, a shuffle distribution was constructed from 250 randomized spike trains. For each randomization, spike times were circularly shifted by a random time offset only across the time epochs of interest. A cell was considered significantly spatially tuned if the Monte-Carlo p-value was less than 0.01.

### Trajectory coding

To characterize trajectory specific coding for individual neurons, a selectivity index was computed as the normalized mean firing rate difference between two trajectories A and B: 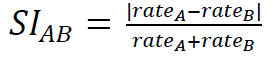. For goal arm specific coding, a selectivity index was computed from the firing rates in the left and right goal arms separately for outbound (*SI_goal|out_*) and inbound (*SI_goal|in_*) journeys. For directional coding, a comparison was made between firing rates in the outbound and inbound journeys (*SI_direction_*). To analyze prospective and retrospective coding, the firing rate on the stem of the Y-maze was compared between outbound journeys towards the left/right goal arms (*SI_prospective_*) and inbound journeys starting from the left/right goal arms (*SI_retrospective_*).

To determine if a selectivity index was higher than expected by chance, a shuffle distribution was constructed by randomly splitting the joint set of trajectories A and B into two new sets A’ and B’ and computing *SI*_*A*′*B*′_ for 250 randomizations, under the null hypothesis that the firing rate is independent of trajectory. *SI*_*AB*_ was considered statistically significant if the Monte-Carlo p-value was smaller than 0.01.

### Neural decoding

Bayesian neural decoding (Davidson et al., 2009; Zhang et al., 1998) was performed to estimate the position and running direction from the spiking activity of isolated lateral septum cells. Position on the track was “linearized” by concatenating the cumulative distances along the three sections of the maze (stem and left/right goal arms). Running direction was defined as a binary variable that represented movements in the inbound and outbound directions. For each cell, a tuning curve λ(*x*, *d*) over linearized position and running direction was computed using compressed kernel density estimation (Sodkomkham et al., 2016) from all times when animals actively ran (speed > 10 cm/s) on stem and goal arms but excluding reward platforms. Given the presence of discontinuities in the linearized position, kernel densities were computed at discrete points on the track using a look-up table with precomputed distances between any two points that conform to the maze topology. For a time bin of duration *Δ* with ***n*** = [*n*_1_, *n*_2_, …, *n*_*ncells*_] spikes for all cells, the posterior probability over linearized position *x* and running direction *d* was computed assuming Poisson firing statistics and a non-informative prior:

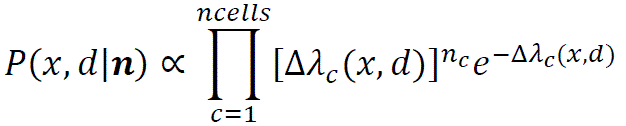

A *K*-fold cross-validation approach was used to evaluate the decoding performance, where *K* represents the total number of outbound and inbound journeys that an animal completed in a session. Thus, tuning curves were computed for all but one journey, and decoding was performed for the left-out journey in *Δ* = 200 ms time bins. This was repeated for all journeys. Posterior probability distributions were summarized by their maximum a posteriori point estimate and decoding performance was quantified as the median distance between estimated and true positions on the maze, and the percentage correct running direction estimates.

To evaluate the dependency of decoding performance on the number of cells included in the analysis, the cross-validated decoding approach was repeated for varying cell sample sizes. For each cell sample size, decoding performance was averaged across 25 different combinations of cells randomly drawn from all available cells. If the number of possible combinations was less than 25, then all combinations were used.

### Neural decoding at theta time scale

For the analysis of spatial and directional representations at the time scale of theta oscillations, neural decoding was performed using the same encoding model as above that includes both LSD and LSI cells. Estimates were computed from spiking activity in *Δ* = 50 ms time bins with 80% overlap. To characterize the temporal dynamics of the decoded spatial estimates, three posterior probability signals were constructed that represent the summed posterior probability in each of the three maze arms. Auto– and cross-correlations were then computed of these signals for four distinct journeys, i.e. when the animals ran towards the choice point (outbound in the stem, or inbound in one of the goal arms) or away from the choice point (outbound in one of the goal arms, or inbound in the stem). On each of the four selected journeys, the animals were either located in the stem or in one of the goal arms, and the corresponding posterior probability signal for that maze arm is referred to as the “local” representation. Similarly, the posterior probability signals for the other maze arms are referred to as “non-local” representations.

To quantify the theta cycle skipping of non-local representations, a cycle skipping index was computed from the auto-correlation, using a modification of the method described above for the spike auto-correlograms. In particular, the correlation values of the first and second theta-associated peaks were measured relative to the minimum correlation in between the peaks, i.e. 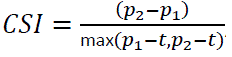, with *t* the minimum correlation value in the 125-250 ms time lag window. Time windows for peaks *p1* and *p2* were adapted to match the observed peaks in the auto-correlograms and set to 80-180 ms and 180-300 ms, respectively.

## Supplemental Figures

**Figure 1 – figure supplement 1.**
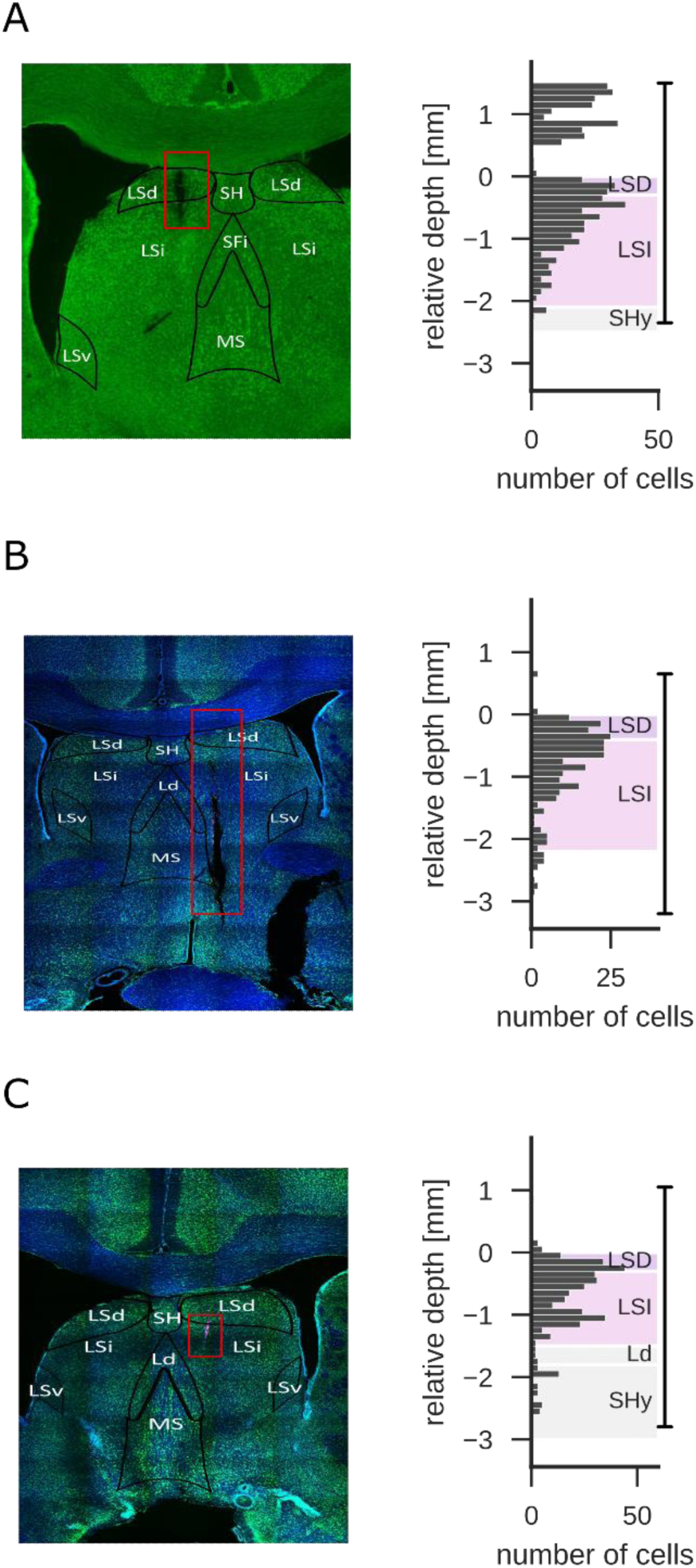
Left: coronal brain slice showing part of the Neuropixels probe trajectory (white arrow) in the lateral septum (animal, A: LS-k-8, B: LS-k-11, C: LS-k-14). Right: the number of recorded cells along the probe shank for one recording session in the same animal. Depth is measured relative to the white matter above the lateral septum. Vertical line at the right indicates the span of the recorded electrodes on the probe.

**Figure 2 – figure supplement 1.**
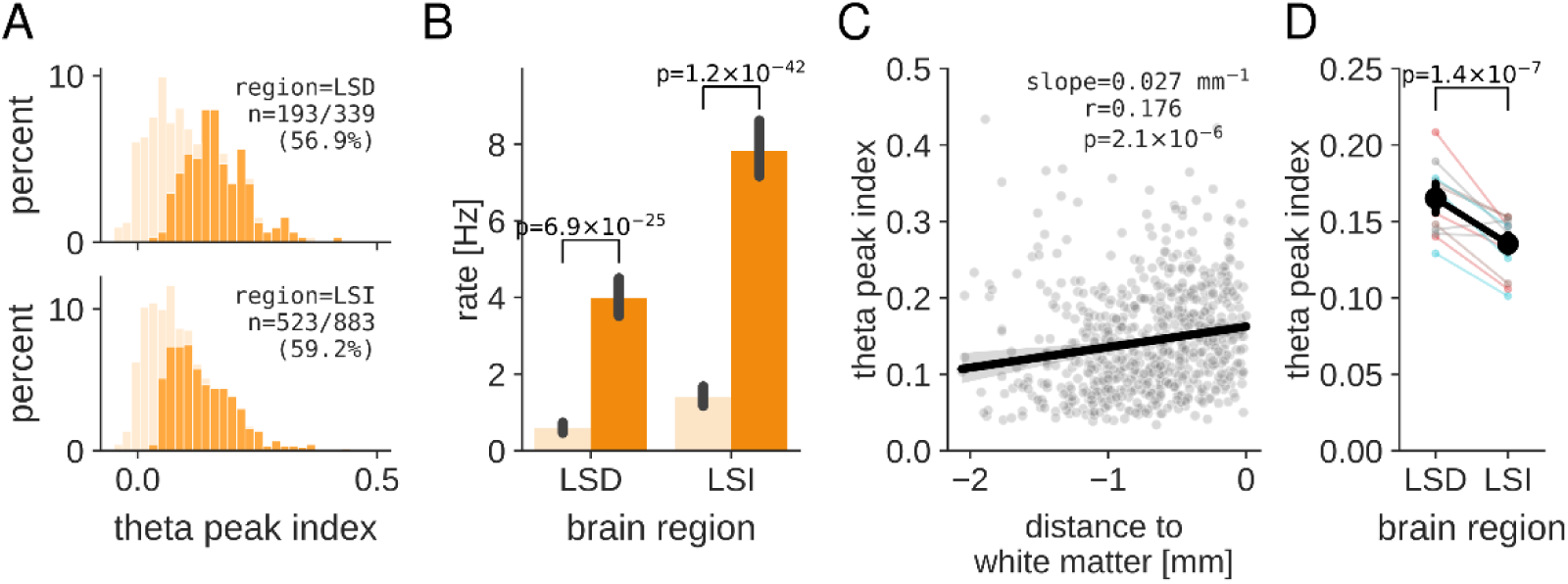
(A) Distribution of theta modulation index of all analyzed neurons in LSD and LSI. Light orange: full distribution of all cells. Dark orange: highlighted part of the distribution that represent cells with significant theta modulation index (Monte-Carlo p-value < 0.01). (B) Mean firing rate of theta rhythmic (dark orange) and non-rhythmic (light orange) cells differ significantly in both lateral septum subregions. Two-sided two-sample t-test, LSD: t(337)=11.18, p=6.9×10^−25^, LSI: t(881)=14.45, p=1.2×10^−42^. (C) Values of theta peak index for theta rhythmic cells increases for cells located closer to the white matter. Dots represent individual cells with significant theta peak index from all sessions and animals. Black line and shaded region represent linear fit and 95% confidence interval (r=0.18, p=2.1×10^−6^). (D) Mean value of theta peak index is significantly higher in LSD as compared to LSI (mean±sem theta peak index, LSD 0.17±0.005, LSI 0.13±0.003; two-sided two-sample t-test, t(714)=5.33, p=1.4×10^−7^). Thin lines represent mean theta peak index for individual sessions, with the line color indicating the animal.

**Figure 3 – figure supplement 1.**
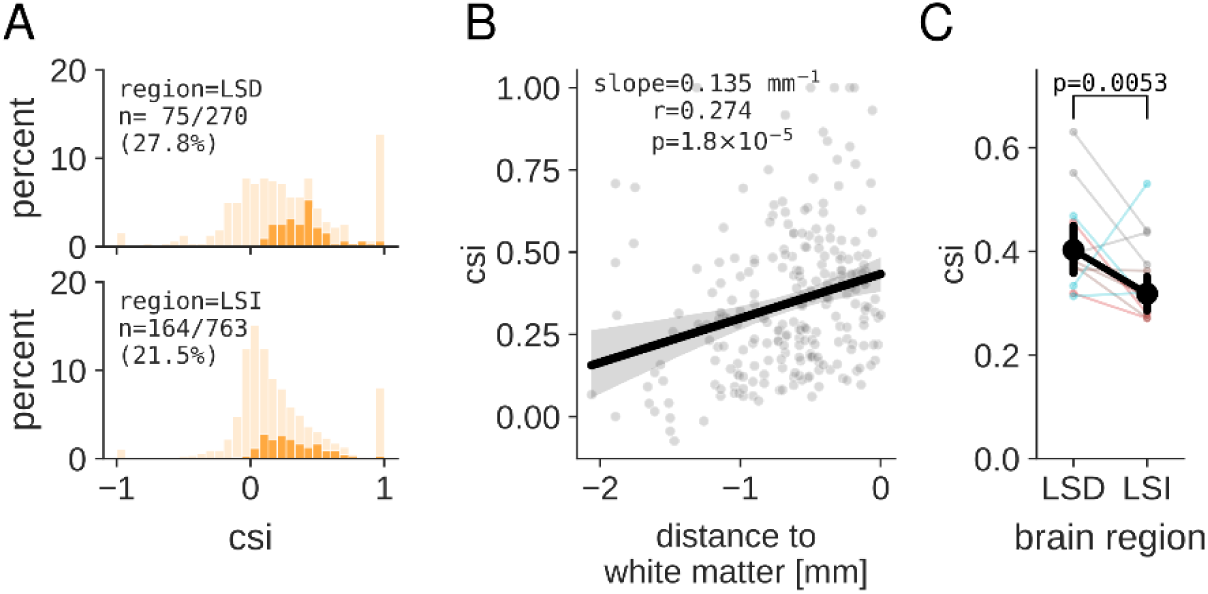
(A) Distribution of CSI values for all analyzed neurons in LSD and LSI. For each cell, the CSI value is taken from the trajectory with the highest z-scored CSI value relative to the shuffle distribution. Light orange: full distribution of all cells. Dark orange: highlighted part of the distribution representing cells with significant CSI value (corrected Monte-Carlo p-value < 0.05). (B) CSI values increase for cells located closer to the white matter. Dots represent individual cells with significant CSI values from all sessions and animals. For each cell, the CSI value is taken from the journey with the highest z-scored CSI value relative to the shuffle distribution. The black line and shaded region represent the linear fit and 95% confidence interval (r=0.27, p=1.8×10^−5^). (C) The mean CSI value of cycle skipping cells is significantly higher in LSD as compared to LSI (mean±sem CSI, LSD 0.40±0.02, LSI 0.32±0.02; two-sided two-sample t-test, t(237)=2.82, p=0.0053). Thin lines represent the mean CSI value for individual sessions, with the line color indicating the animal.

**Figure 3 – figure supplement 2.**
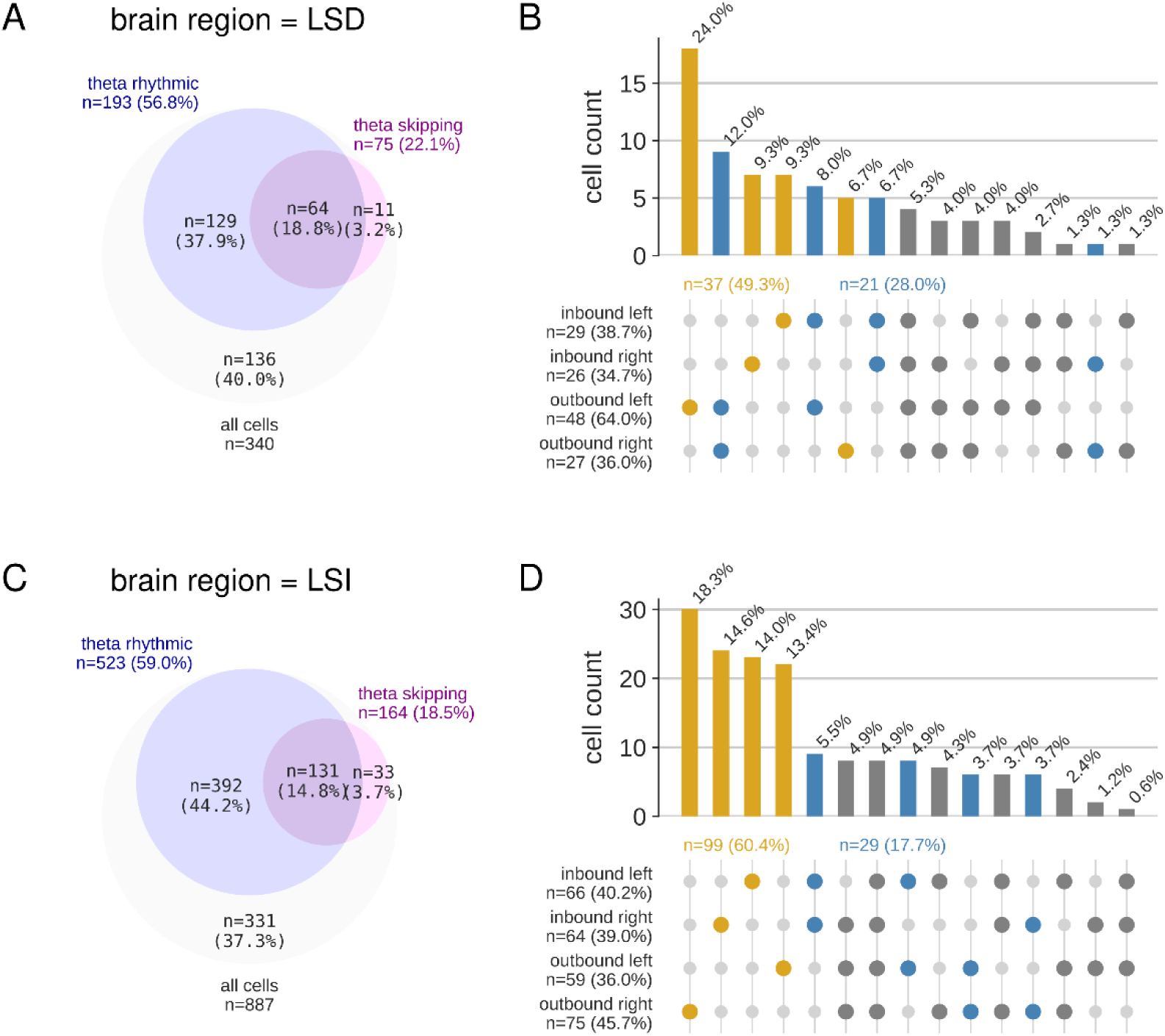
(A) Venn diagram showing overlap of cell populations with significant theta rhythmicity and theta cycle skipping in LSD. (B) Histogram of the number of cells in LSD with a significant cycle skipping index for all possible journey combinations. Note that for most cells (49.3%; yellow) cycle skipping occurs only on a single journey type. For another population of cells (28.0%; blue), cycle skipping occurs on outbound, inbound, left, or right journeys. (C) Venn diagram showing overlap of cell populations with significant theta rhythmicity and theta cycle skipping in LSI. (D) Histogram of the number of cells in LSI with a significant cycle skipping index for all possible journey combinations. Note that for most cells (60.4%; yellow) cycle skipping occurs only on a single journey type. For another population of cells (17.7%; blue), cycle skipping occurs on outbound, inbound, left, or right journeys.

**Figure 5 – figure supplement 1.**
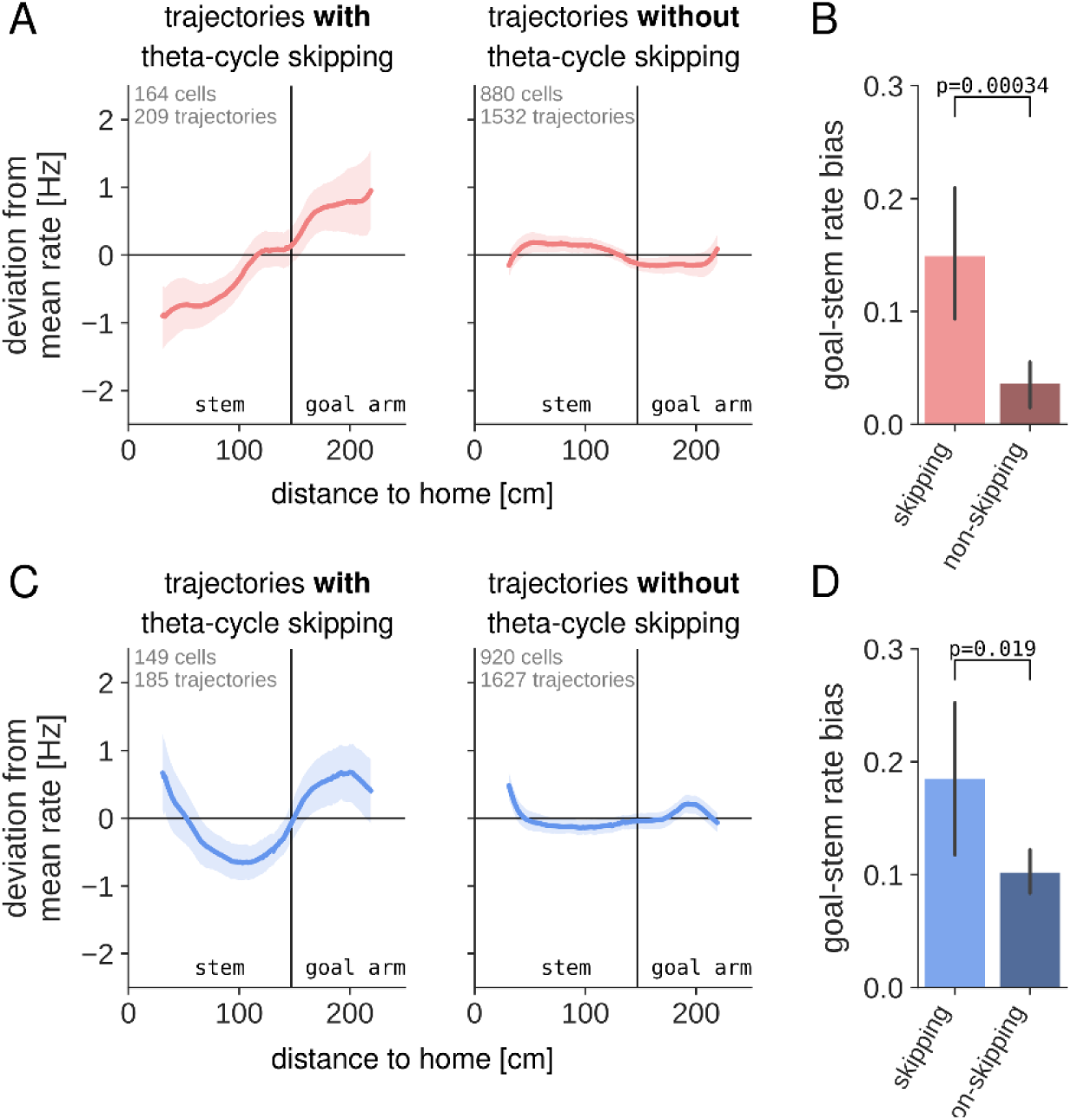
(A) Left: lateral septum cells that exhibit theta cycle skipping on outbound trajectories increase their firing rate from stem to goal arm. The trajectory-specific (i.e., outbound left and outbound right) spatial tuning curves were mean-corrected for each cell. The solid line represents the average rate deviation for all cell-outbound trajectory pairs with significant theta cycle skipping. The shaded area represents the 95% CI. Right: No increase in firing rate was present for cell-outbound trajectory combinations without theta cycle skipping. (B) Quantification of the goal arm rate bias across all cell-outbound trajectory pairs with or without significant theta cycle skipping. Welch’s independent samples t-test: t(260.3)=3.63, p=0.00034. (C) The same analysis as in (A), but for inbound trajectories. (D) Quantification of the goal arm rate bias across all cell-inbound trajectory pairs with or without significant theta cycle skipping. Welch’s independent samples t-test: t(219.1)=2.36, p=0.019.

**Figure 6 – figure supplement 1.**
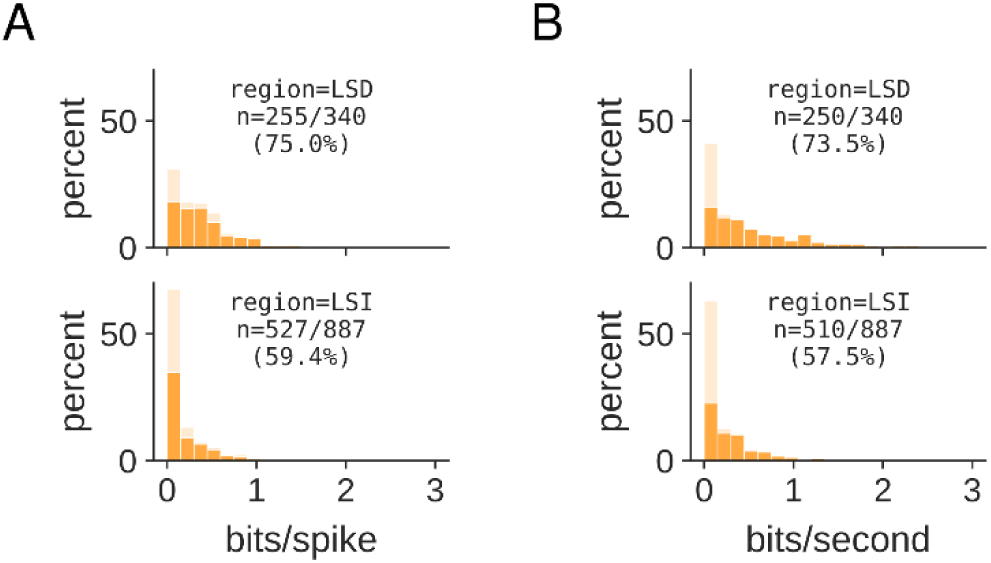
Distribution of spatial information in bits/spike (A) and bits/second (B) of all analyzed neurons in LSD (top) and LSI (bottom). Light orange: full distribution of all cells. Dark orange: highlighted part of the distribution that represents cells with significant spatial information (Monte-Carlo p-value < 0.01). Overall mean±sem of significant cell population, spatial information in bits/spike: LSD 0.42±0.02, LSI 0.21±0.01; spatial information in bits/second: LSD 0.69±0.05, LSI 0.37±0.02. Welch’s independent t-test comparing spatial information of significant cell populations in LSD and LSI, bits/spike: t(358.5)=8.17, p=5.3×10^−15^; bits/second: t(330.0)=6.38, p=6.0×10^−10^.

**Figure 6 – figure supplement 2.**
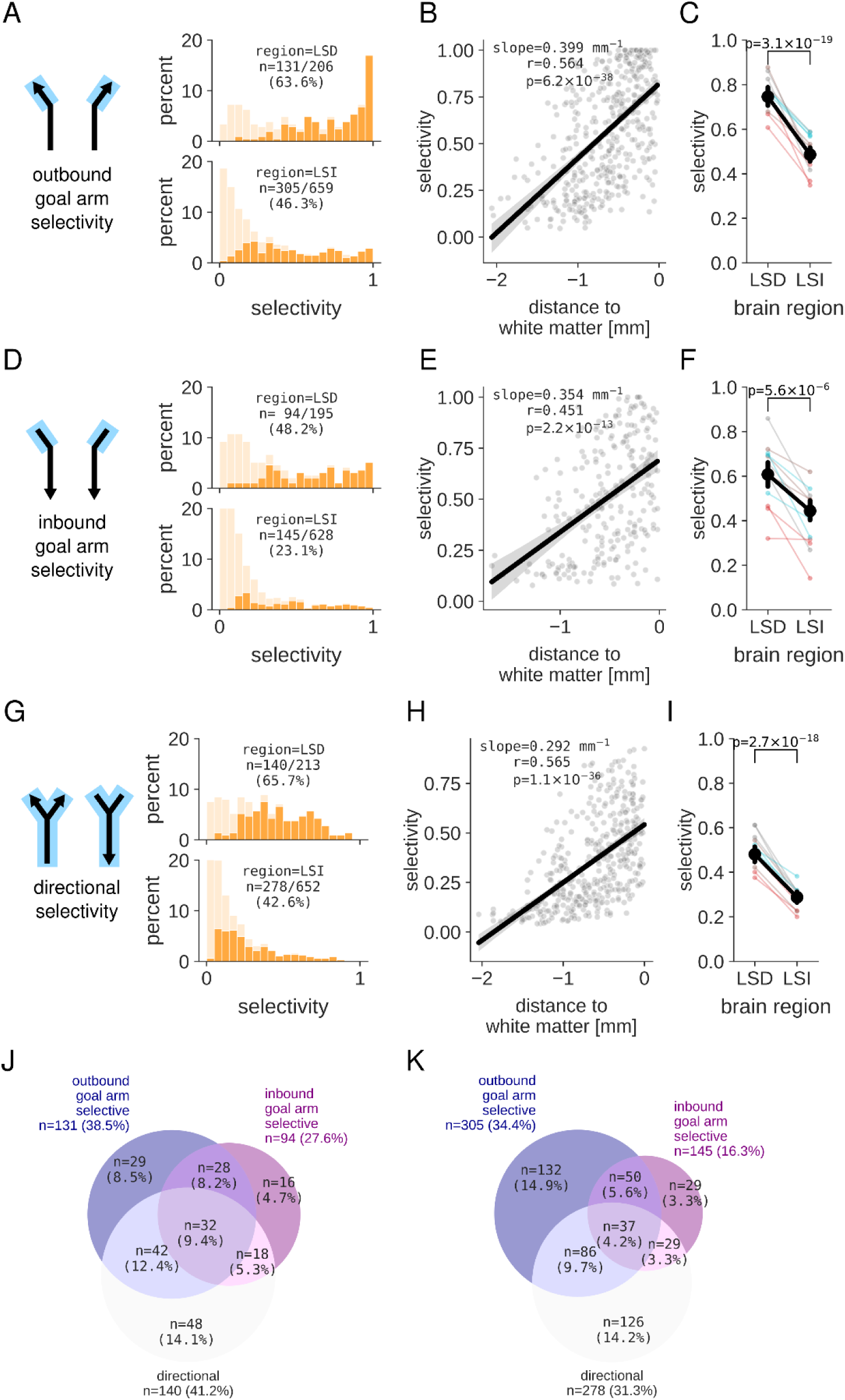
(A,B,C) Distribution of outbound goal arm (A), inbound goal arm (B) and directional (C) selectivity of all analyzed neurons in LSD and LSI. Light orange: full distribution of all cells. Dark orange: highlighted part of the distribution that represent cells with significant selectivity (Monte-Carlo p-value < 0.01). (D,E,F) Values of outbound goal arm (D), inbound goal arm (E) and directional (F) selectivity index increases for cells located closer to the white matter. Dots represent individual cells with significant selectivity index from all sessions and animals. Black line and shaded region represent linear fit and 95% confidence interval. (G,H,I) Mean value of outbound goal arm (G), inbound goal arm (H) and directional (I) selectivity index is significantly higher in LSD as compared to LSI (mean±sem selectivity, outbound goal arm: LSD 0.75±0.02, LSI 0.49±0.02, two-sided two-sample t-test, t(434)=9.40, p=3.1×10^−19^; inbound goal arm: LSD 0.61±0.03, LSI 0.44±0.02, two-sided two-sample t-test, t(237)=4.65, p=5.6×10^−6^; directional: LSD 0.48±0.02, LSI 0.29±0.01, two-sided two-sample t-test, t(416)=9.14, p=2.7×10^−18^). (J,K) Overlap between goal arm and directional selectivity for cells in LSD (J) and LSI (K).

**Figure 6 – figure supplement 3.**
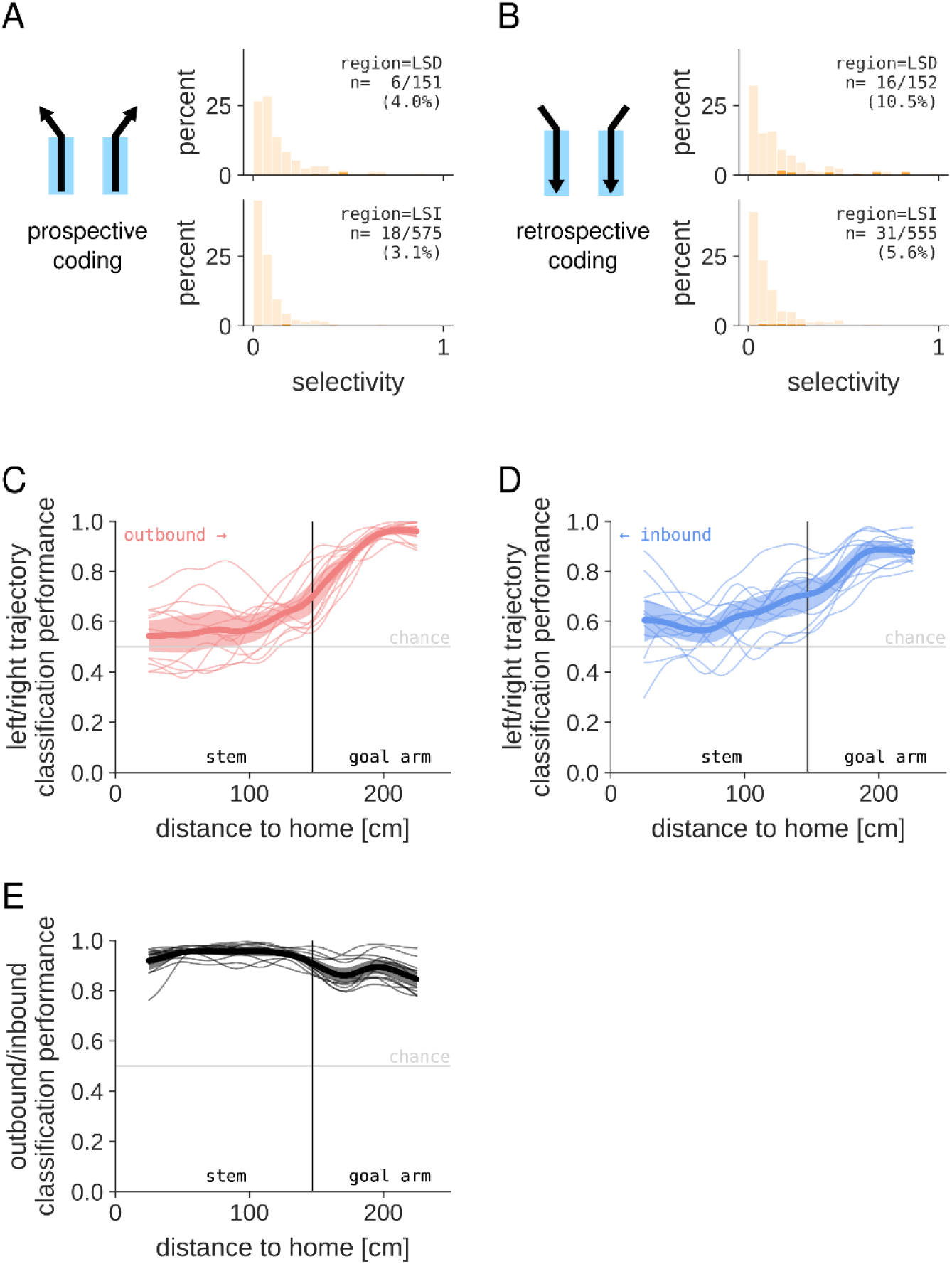
(A) Distribution of prospective coding of all analyzed neurons in LSD and LSI. Light orange: full distribution of all cells. Dark orange: highlighted part of the distribution representing cells with significant prospective coding (Monte-Carlo p-value < 0.01). (B) Distribution of retrospective coding of all analyzed neurons in LSD and LSI. Light orange: full distribution of all cells. Dark orange: highlighted part of the distribution representing cells with significant retrospective coding (Monte-Carlo p-value < 0.01). (C) Binary classification of left/right outbound trajectory as a function location on the maze. Thin lines: individual sessions; thick line: mean across sessions; shaded region: 95% CI. (D) Binary classification of left/right inbound trajectory as a function location on the maze. Thin lines: individual sessions; thick line: mean across sessions; shaded region: 95% CI. (E) Binary classification of run direction (outbound or inbound) as a function of location on the maze. Thin lines: individual sessions; thick line” mean across sessions; shaded region: 95% CI.

**Figure 9 – figure supplement 1.**
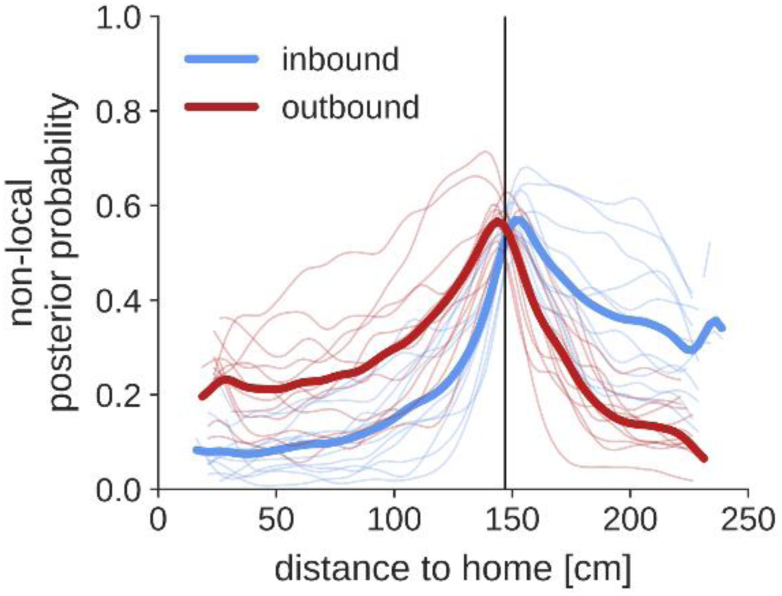
Average posterior probability following theta time scale decoding that is assigned to non-local maze arms (i.e., the arms where the animal is not currently located). Data for the goal arms is combined and position is expressed as distance to home. Vertical line indicates the choice point. Thin lines represent data for individual sessions; thick lines represent average across sessions.

**Figure 10 – figure supplement 1.**
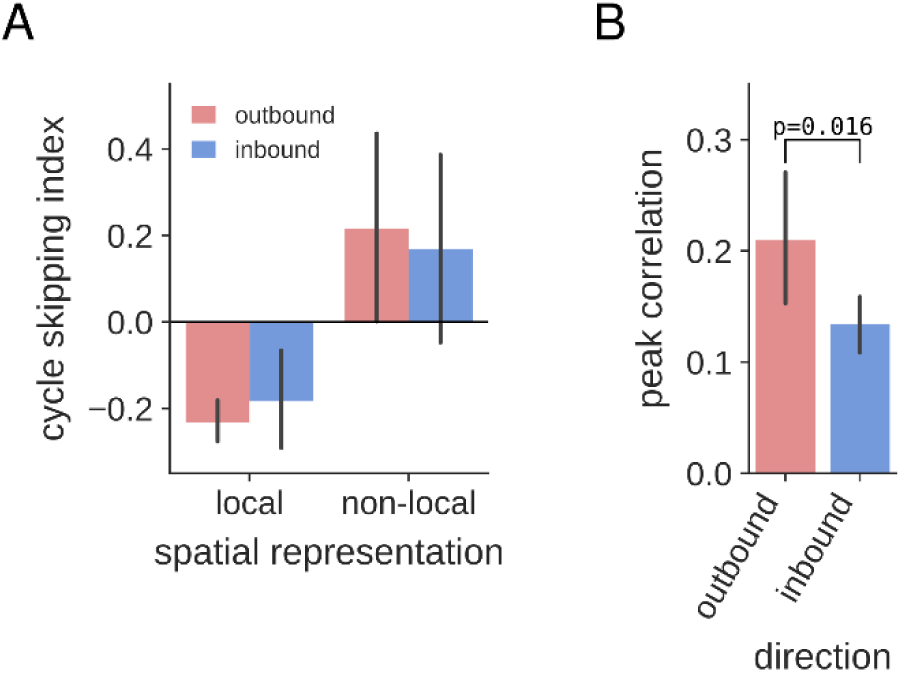
(A) Cycle skipping index computed from the auto-correlations of local and non-local spatial representations, separately for outbound and inbound trajectories. Two-way ANOVA for the effects of locality and run direction on cycle skipping index: no significant interaction between locality and run direction, F(1,44)=0.30, p=0.59; a significant main effect for locality, F(1,44)=20.77, p=4.1×10^−5^; no significant main effect for run direction, F(1,44)=7.7×10^−5^, p=0.99). (B) Peak cross-correlation between the two non-local spatial representations for outbound and inbound trajectories. Paired samples t-test: t(11)=2.83, p=0.016.

**Figure 10 – figure supplement 2.**
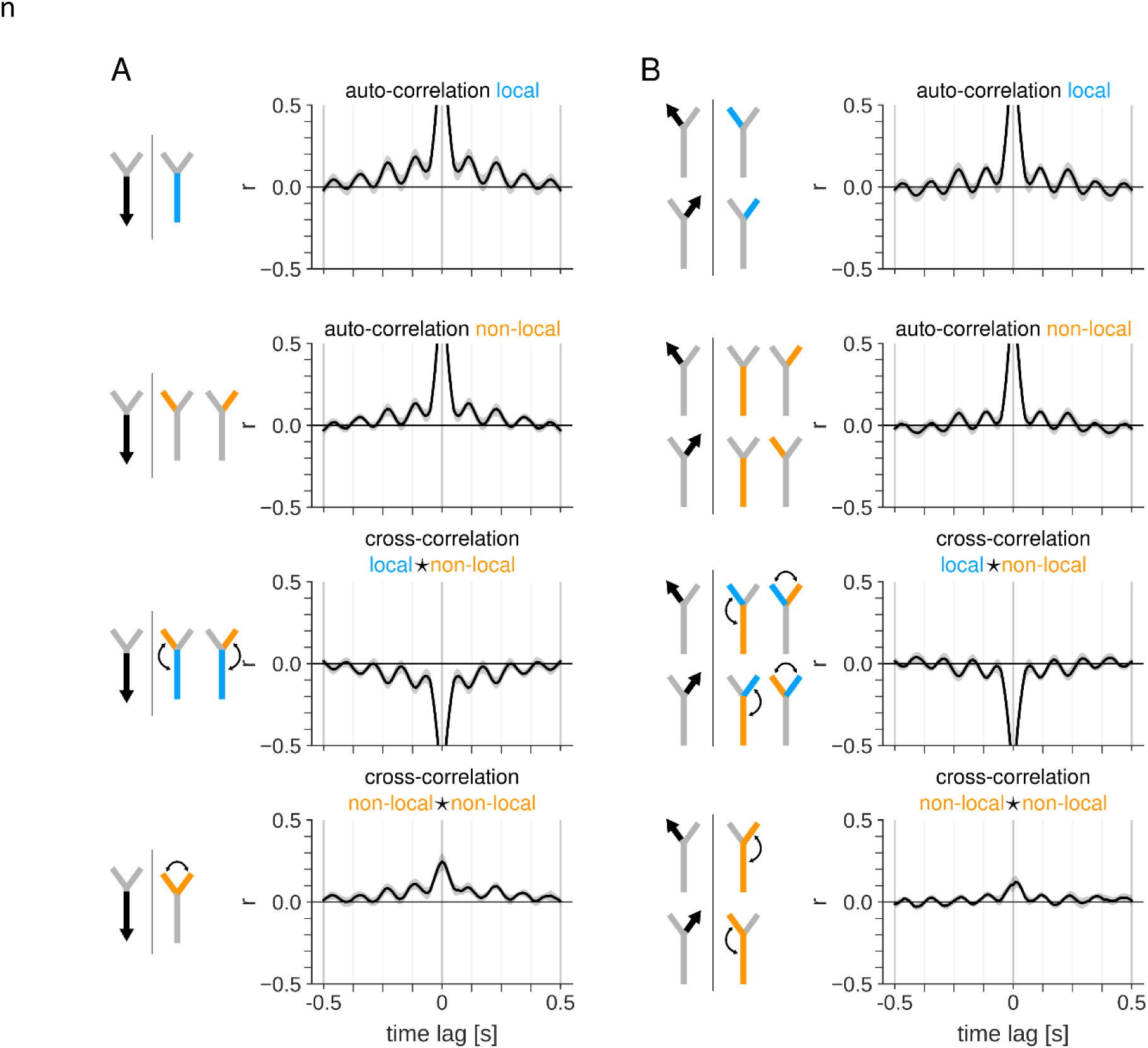
(A) Auto– and cross-correlation of posterior probability time courses for the three maze sections when the animal is running along the stem in the inbound direction towards home. Correlations are computed as the Pearson correlation coefficient at varying time lags. For each plot, drawings at the left show the animal’s behavior (black arrow) and the maze sections for which correlations are computed (color indicates whether the highlighted maze section is local (blue) or non-local (orange) relative to the animal’s position on the track). Top: auto-correlation of local representations in the stem. Second from top: auto-correlations of non-local representations in the two goal arms. Third from top: cross-correlations between local and non-local representations. Bottom: cross-correlation between non-local representations in the two goal arms. (B) Same as (A), but for times when the animal is running along one of the goal arms in the outbound direction towards the reward platform. Equivalent correlations for the two goal arms are computed jointly.

**Figure 11 – figure supplement 1.**
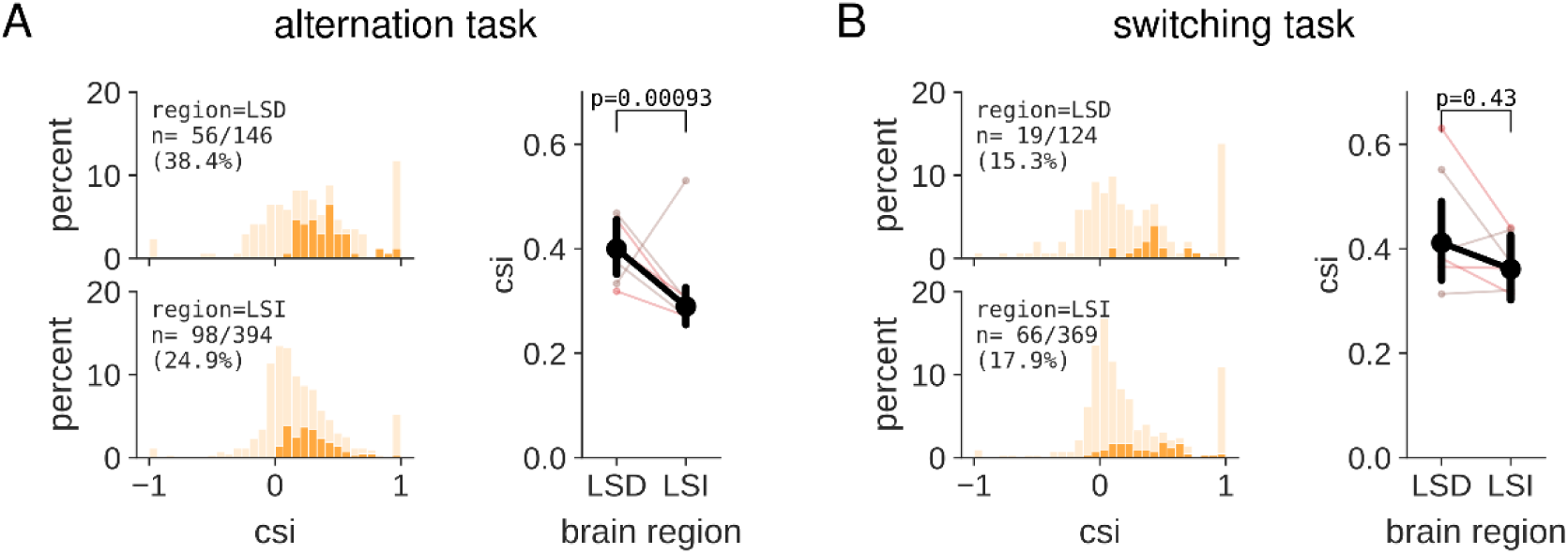
(A) Left: distribution of CSI values for LSD (top) and LSI (bottom) neurons in sessions in which rats performed the alternation task. For each cell, the CSI value is taken from the trajectory with the highest z-scored CSI value relative to the shuffle distribution. Light orange: full distribution of all cells. Dark orange: highlighted part of the distribution that represent cells with significant CSI value (corrected Monte-Carlo p-value < 0.05). Right: mean CSI value is significantly higher in LSD as compared to LSI (mean±sem CSI, LSD 0.40±0.03, LSI 0.29±0.02; two-sided two-sample t-test, t(152)=3.38, p=0.00093). Thin lines represent mean CSI value for individual sessions, with the line color indicating the animal. (B) Same as (A), but for sessions in which animals performed the switching task. Mean CSI value (right) is not different between LSD and LSI (mean±sem CSI, LSD 0.41±0.04, LSI 0.36±0.03; two-sided two-sample t-test, t(83)=0.79, p=0.43).

**Figure 11 – figure supplement 2.**
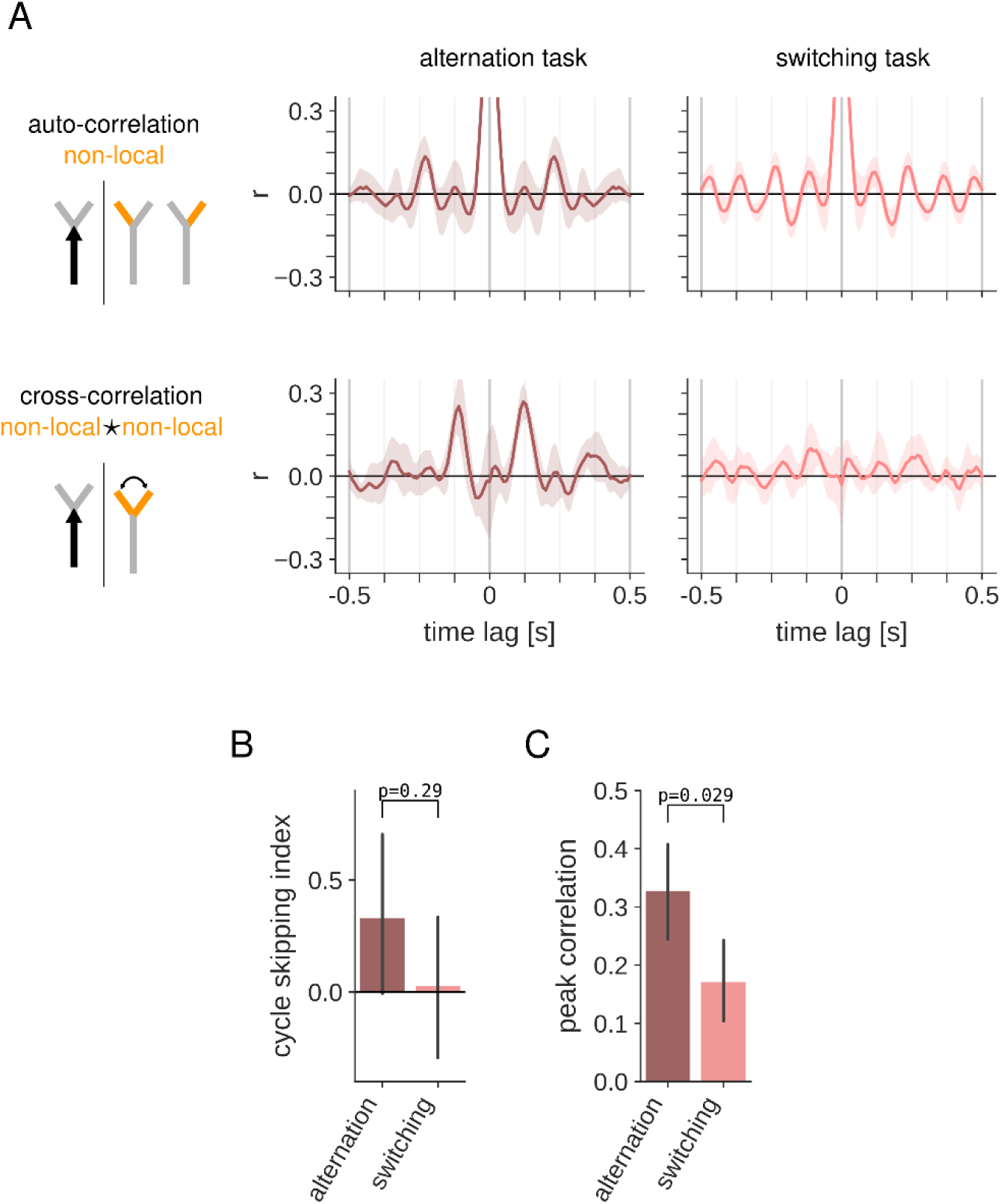
(A) Auto– and cross-correlations of the non-local spatial representations in the first five outbound trajectories in the alternation task (left) and switching task (right). (B) Task comparison of the cycle skipping index computed from the auto-correlation of non-local spatial representations in the first five outbound trajectories of each session. Independent samples t-test: t(10)=1.12, p=0.29. (C) Task comparison of the peak cross-correlation of non-local spatial representations in the first five outbound trajectories of each session. Independent samples t-test: t(10)=2.54, p=0.029.

**Figure 11 – figure supplement 3.**
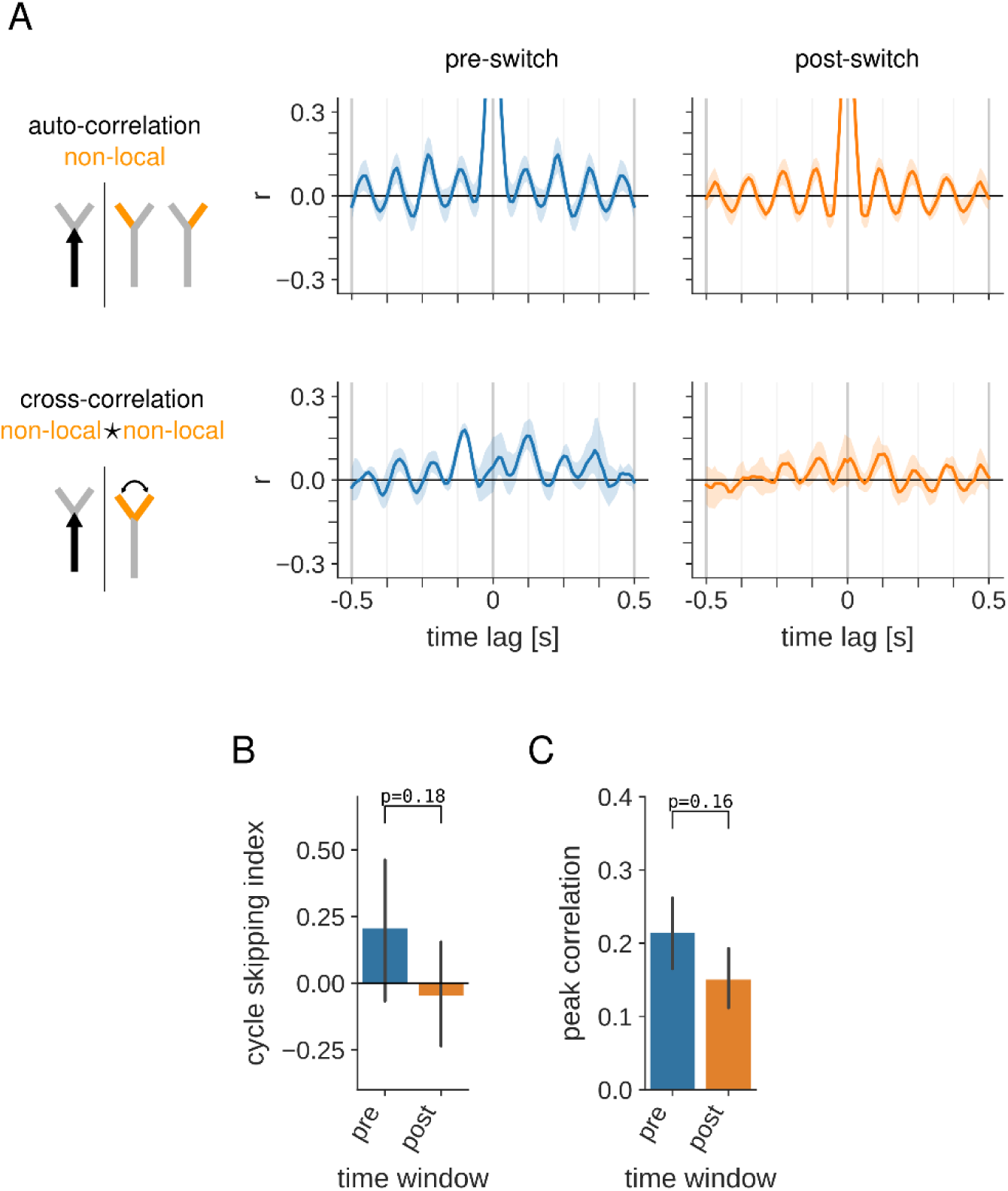
(A) Auto– and cross-correlations of the non-local spatial representations in the five trials before (left) and after (right) the change in reward contingency in switching task sessions. Only outbound trajectories are included in the analysis. (B) Pre– vs post-switch comparison of the cycle skipping index computed from the auto-correlation of non-local spatial representations in outbound trajectories. Paired samples t-test: t(5)=1.57, p=0.018. (C) Pre– vs post-switch comparison of the peak cross-correlation of non-local spatial representations in outbound trajectories. Paired samples t-test: t(5)=1.64, p=0.016.

**Figure 11 – table supplement 1.**
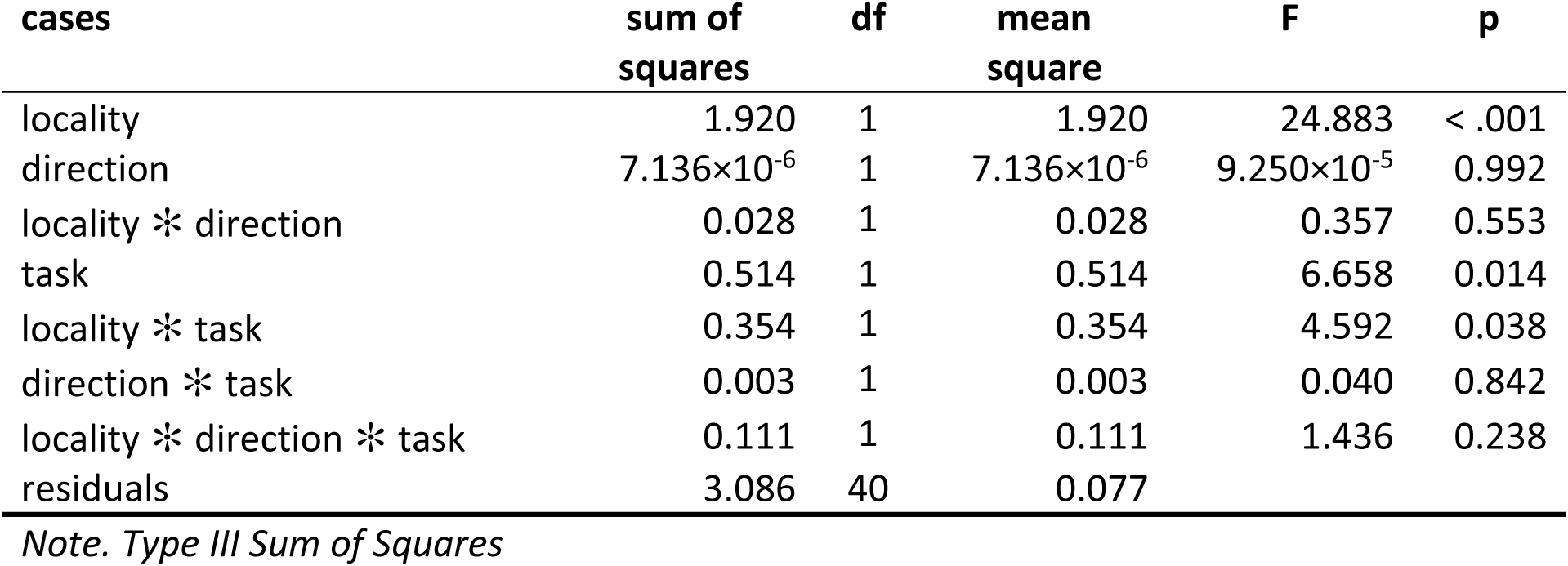
Three-way ANOVA for the effect of locality, direction, and task on the cycle skipping index of decoded spatial representations.

**Figure 11 – table supplement 2.**
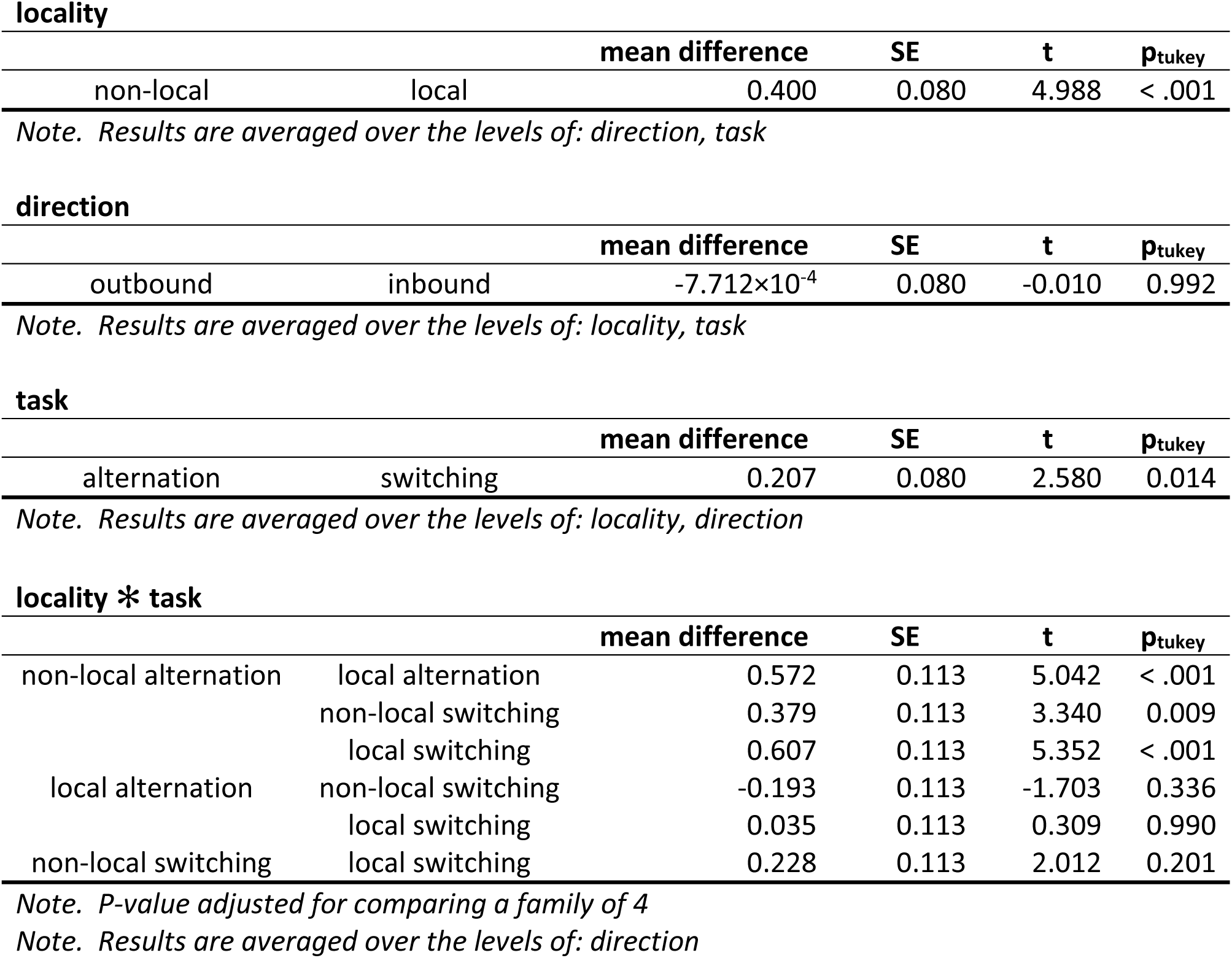
Post-hoc comparisons for three-way ANOVA in Figure 11 – table supplement 1.

**Figure 11 – table supplement 3.**
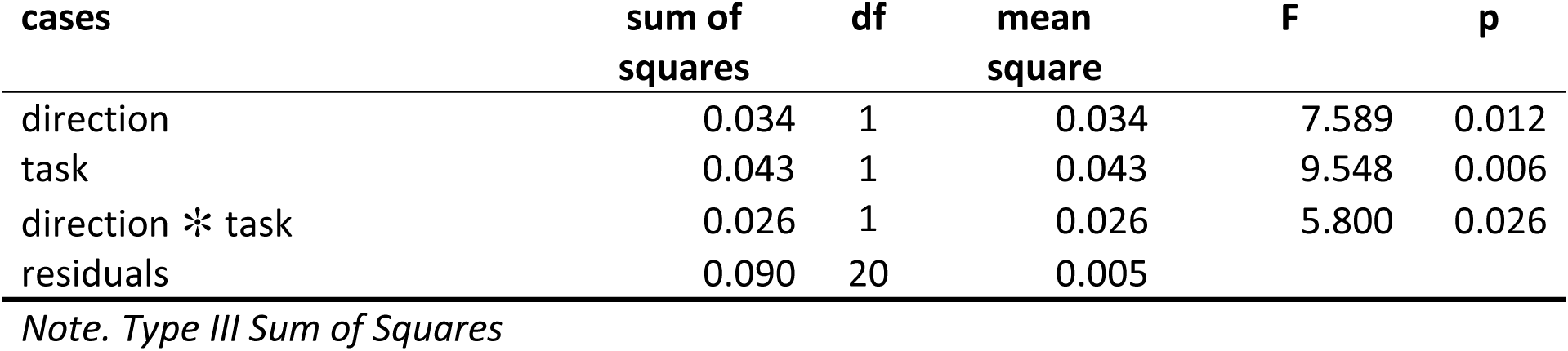
Two-way ANOVA for the effect of direction and task on the peak cross-correlation of non-local spatial representations.

**Figure 11 – table supplement 4.**
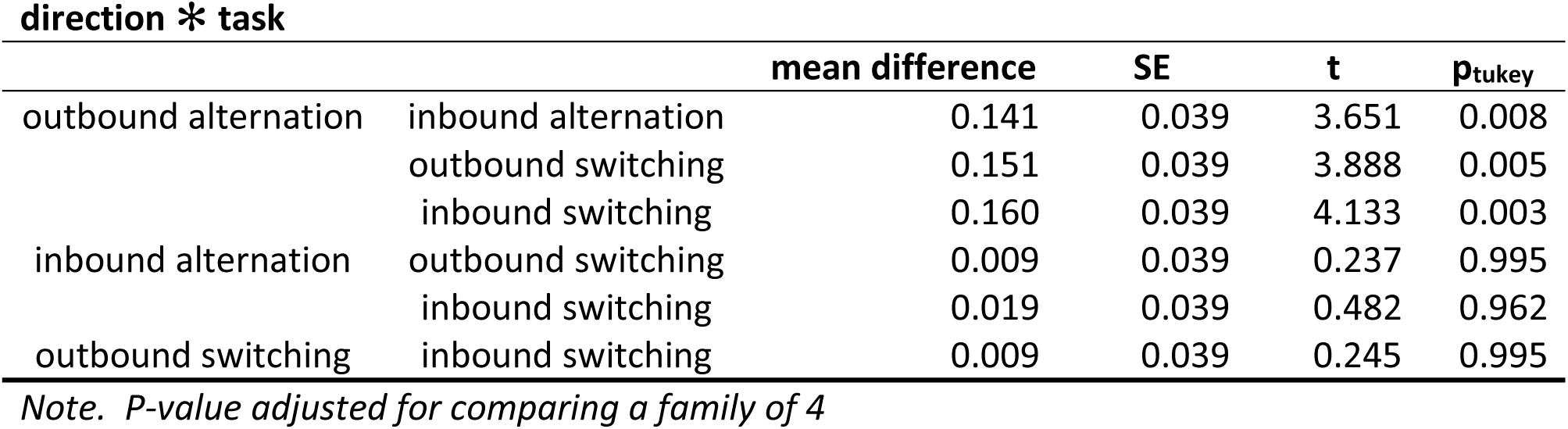
Post-hoc comparisons for two-way ANOVA in Figure 11 – table supplement 3.

## Author contributions

**Katarzyna Bzymek**: conceptualization, formal analysis, investigation, writing-original draft, writing – review & editing, visualization

**Fabian Kloosterman**: conceptualization, formal analysis, writing-original draft, writing – review & editing, visualization, supervision, funding acquisition

## Competing interests

No competing interests declared.

## Acknowledgments

This work was supported by grants to F.K. from The Research Foundation – Flanders (FWO), grant numbers G0D7516N and G077321N.

## Data Availability Statement

The data, Python code and notebooks generated in this study that were used to create all figures and to compute statistics have been deposited in a public Open Science Framework repository (https://osf.io/djaem/).

## Notes

https://osf.io/djaem/

### Competing Interest Statement

The authors have declared no competing interest.

### Summary of Updates

This revised manuscript contains updated quantification of theta cycle skipping, adds new statistical comparisons of the difference between the two behavioral tasks, and includes general improvements to the text and figures.

